# Exploration of cell development pathways through high dimensional single cell analysis in trajectory space

**DOI:** 10.1101/336313

**Authors:** Denis Dermadi, Michael Bscheider, Kristina Bjegovic, Nicole H. Lazarus, Agata Szade, Husein Hadeiba, Eugene C. Butcher

**Author notes:** shared correspondence. shared authors (Denis Dermadi, Michael Bscheider). Author contributions: DD wrote the algorithm, supervised computational analyses, and interpreted intestinal data; KB wrote parts of the algorithm; DD & HH designed and interpreted the thymus study; MB & NHL performed and DD & MB analyzed the tonsil B cell study; AS prepared human tissues; DD, MB, ECB wrote the manuscript; HH provided advice; ECB conceived the trajectory space concept and supervised the project.

## Abstract

High-dimensional single cell profiling coupled with computational modeling is emerging as a powerful means to elucidate developmental sequences and define genetic programs directing cell lineages. Here we introduce tSpace, an algorithm based on the concept of “trajectory space”, in which cells are defined by their distance along nearest neighbor pathways to every other cell in a population. tSpace outputs a dense matrix of cell-to-cell distances that quantitatively reflect the extent of phenotypic change along developmental paths (developmental distances). Graphical mapping of cells in trajectory space allows unsupervised reconstruction and straightforward exploration of complex developmental sequences. tSpace is robust, scalable, and implements a global approach that attempts to preserve both local and distant relationships in developmental pathways. Applied to high dimensional flow and mass cytometry data, the method faithfully reconstructs known pathways of thymic T cell development and provides novel insights into regulation of tonsillar B cell development and trafficking. Applied to single cell transcriptomic data, the method unfolds complex developmental sequences, recapitulates pathways leading from intestinal stem cells to specialized epithelial phenotypes more faithfully than existing algorithms, and reveals genetic programs that correlate with fate decisions. tSpace profiling of complex populations in high-dimensional trajectory space is well suited for hypothesis generation in developing cell systems.

---

Precursor cells give rise to differentiated progeny through complex developmental pathways. Single cell technologies hold the promise of elucidating the developmental progression and defining underlying transcriptomic drivers and modulators. Mass cytometry (CyTOF) and single cell RNA-seq (scRNAseq) can capture a high-dimensional profile of a “cellular snapshot” within analyzed tissue that contains all developing, renewing and differentiated cell populations. High-dimensional profiles of cells can then be computationally aligned to reveal developmental relationships.

Here we show that developmental pathways can be reconstructed from single cell profiles by analyzing cells in “trajectory space”, in which each cell is represented by a profile or vector of its relative distances along nearest neighbor pathways to every other cell. The concept is illustrated in Fig. 1a, with a schematic example of several cells derived from cell A and analyzed with two phenotypic markers. Cells H and E are phenotypically similar but arise from different developmental sequences and thus are developmentally distant. A dense matrix of cell-to-cell distances along the developmental pathways is constructed, which when visualized with standard dimensionality reduction tools [e.g. principal component analysis (PCA)] can be used to explore cell relationships in this novel trajectory space. As illustrated, the method reconstitutes the correct branching developmental sequences of cells in the simple example.

**Figure 1.**
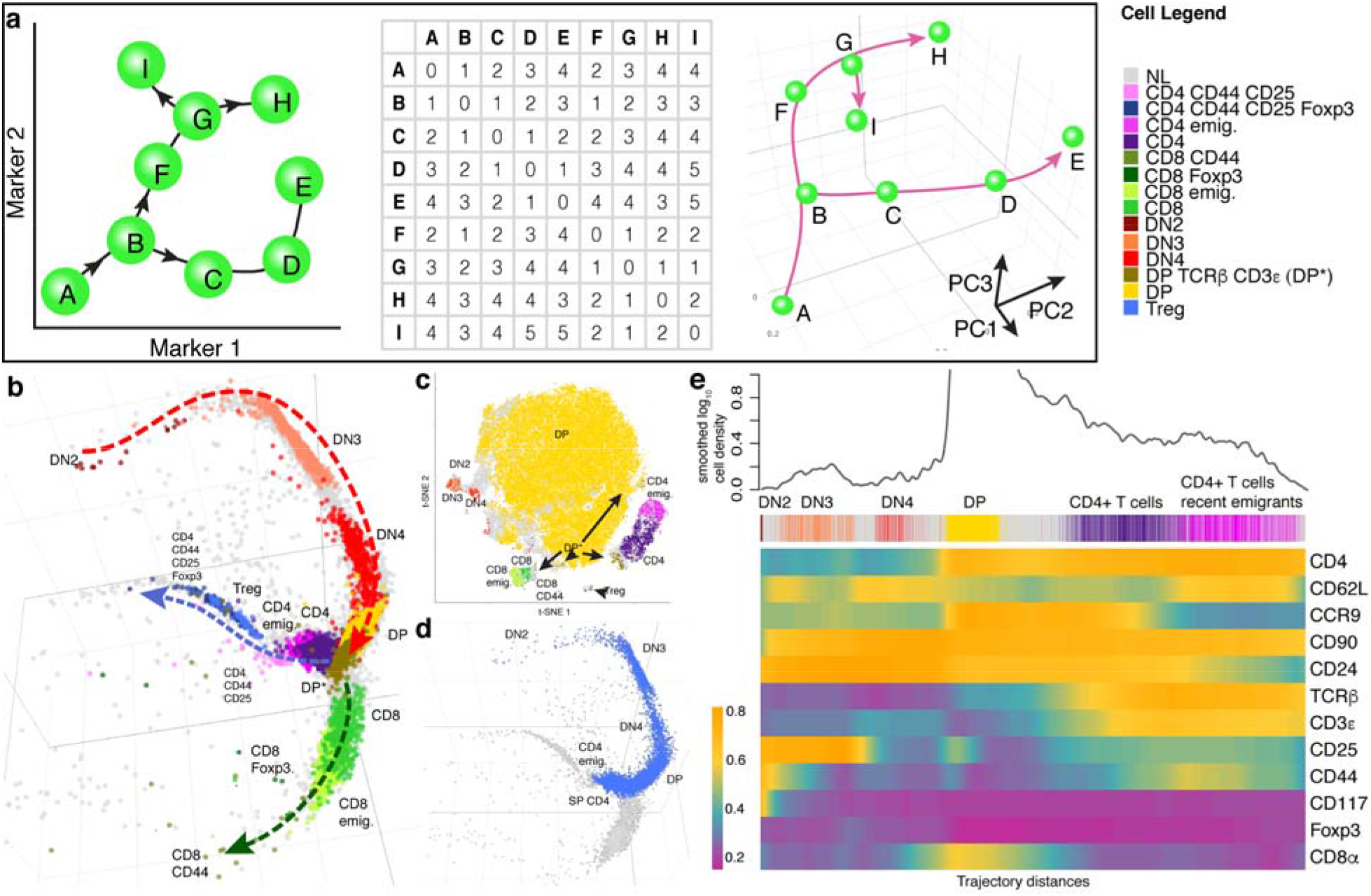
tSpace concept and application on thymic T cells: tSpace reveals developmental trajectories and recovers expression patterns of markers of T cell differentiation. **a** A schematic example illustrates the concept of trajectory space. The “cells” are marked with the letters (A-I) and their developmental sequences with arrows. A matrix of cell-to-cell distances along developmental paths is created (each cell is one unit from its nearest neighbor). Visualization of cell positions in this ‘trajectory space’, here using PCA, recapitulates the branches. Note that E and H, although similar phenotypically, are most distant in trajectory space reflecting their developmental pathways. **b** Unsupervised tSpace analysis of thymic mouse thymocytes accurately recapitulates thymic T cell development. **c** t-SNE of thymic T cells defines clusters but not developmental relationships. **d** Isolated trajectory from DN2 precursors to CD4 thymic emigrants. **e** Smoothed expressions of measured markers along isolated trajectory (shown in d) reveals patterns of protein regulation during T cell differentiation. The identities of manually defined cell subsets as well as cell density along the isolated trajectory are shown for reference above the heatmap. DN - double negative T cells; CD4 emig. - CD4^+^ T cell poised emigrants; CD8 emig. - CD8^+^ T cell poised emigrants.

To implement the concept, we developed a tSpace algorithm. Its application to single cell datasets relies on the assumptions that (i) developmental processes are gradual, (ii) all developmental stages are represented in the data and (iii) markers used to profile cells are regulated and sufficiently informative to distinguish different developmental pathways. Starting with cell profiles (phenotypes), tSpace identifies the K nearest neighbors of every cell, constructs a nearest neighbor (NN) graph that provides connections to all cells in the dataset; calculates distances from each cell to every other cell in the population along NN connections; and exports a dense matrix of N × T dimensions (number of cells N × number of calculated “trajectories” T, vectors of cell-to-cell distances within the manifold). Unlike other tools, tSpace determines the distances within the KNN graph using Wanderlust^1^, an algorithm that takes advantage of subgraphs and waypoints and implements a weighting scheme to reduce “short-circuits” in selecting optimal paths. The Wanderlust algorithm has been described in detail^1^. It significantly improves the definition of branching pathways even in simple flow cytometry datasets (Fig. S1). We outline the effects of varying user-defined Wanderlust parameters on tSpace in the Supplementary Methods (Fig. S2). tSpace detection of developmental relationships is robust over a range of input parameters, allowing implementation of default settings that work well in different applications. The tSpace output provides principal component and UMAP embedding of cells in trajectory space, suitable for visualization and biological exploration of developmental pathways. The cell-to-cell distances, when exported, provide quantitative measures of phenotype change within the manifold based on user selected metrics, useful for ‘pseudotime’ ordering and analysis of e.g. gene/protein expression changes along isolated linear developmental sequences. The use of unified metric-defined distances enables comparison of protein or gene expression dynamics along different trajectories on a common axis.

For samples with large cell numbers (N), tSpace has the option of calculating fewer trajectories, but it is important that these trajectories start from cells well distributed throughout phenotypical space. K-means clustering identifies groups of cells that are well distributed within phenotypic space, and we calculate trajectories from one cell from each such cluster. The clusters are not used for further analysis. As illustration of this feature, tSpace accurately recapitulates simple branching developmental paths from as few as 25 – 100 trajectories (Fig. S1).

To evaluate the algorithm, we applied tSpace to developing populations of lymphocytes analyzed by flow or mass cytometry, and of small intestinal epithelial cells analyzed by scRNAseq.

T cell development in the thymus is well established and allows validation of tSpace in a defined system. We generated flow cytometric profiles of mouse thymocytes using a panel of 13 antibodies (Supplementary Table 1). Our panel detects early T cell populations (so-called ‘double negative’ populations DN1-DN4, which lack CD4 and CD8 and are distinguished by CD44 and CD25 expression), double positive (DP) CD4^+^CD8^+^ cells, and CD4 or CD8 single positive (SP) T cells including poised thymic emigrant phenotype cells, regulatory T cells (CD4^+^, CD25^+^, Foxp3^+^) and a small fraction of SP T cells expressing CD44, an activation and memory marker. We manually gated on these subsets and labeled them (Fig. S3)^2^. Unsupervised tSpace analysis reveals the expected bifurcation of CD4 vs CD8 lineages from the dominant DP population in thymopoiesis and correctly positions T cells from early (DN2) to mature thymic emigrant phenotype T cells in known developmental relationships (Fig. 1b). DN1 cells were not present in the dataset. In addition to the expected major bifurcation of CD4 vs CD8 cells arising from the dominant DP pool, the analysis reveals branching of regulatory T cells (Foxp3^+^) from the SP CD4 stage of CD4 branch. In contrast to methods based on or using clustering for visualization (e.g. PAGA, SPADE, p-Creode, see comparisons in supplementary materials), tSpace highlights a developmental continuum of cells allowing exploration of intermediate populations. For example, tSpace visualizes DP cells in transition to the more mature SP CD4 and CD8 T cells. The transitional cells co-express CD4 and CD8 but some have upregulated TCRβ and CD3ε, a characteristic of positively selected cells^3^. Conventional clustering, based on measured markers using t-SNE, identifies the major subsets but does not clarify developmental relationships (Fig. 1c).

The tSpace output allows evaluation of expression of markers along developmental paths. To illustrate this for CD4 cell differentiation, we manually gated on cells along the path from DN2 cell to CD4 thymic poised emigrants (Fig. 1d), identified and averaged trajectories in the exported tSpace matrix (Methods) that started from early DN2 cells, and displayed marker expression along their trajectory distance from DN2 cells in a heatmap (Fig. 1e). The results capture regulation of marker proteins as cells progress towards maturity, recapitulating known phenotypic progression of thymic T cell development and highlighting details of transitional states. For example, protein expression trends confirm upregulation of the chemokine receptor CCR9 in DN3 cells but reveal notably stronger expression in DN4-DP transitioning cells. CCR9 binds CCL25 expressed by thymic epithelial cells and promotes T cell cortical positioning^4^.

Single cell analyses hold the potential to provide insights into patterns of cell development in settings not accessible to experimental manipulation, as in the human. We applied tSpace to the development of B cells in human tonsils. Naïve (IgD^+^) B cell differentiation towards Immunoglobulin A or G (IgA, IgG) class-switched memory or plasma cells has been investigated. However, the sequence of class switch- and fate-determining decision points and trafficking receptor induction remain poorly defined^5,6^. We used a panel of mass labeled antibodies that detects ~25 markers of B cell subsets and maturation (Supplementary Table 2) to stain human tonsil lymphocytes. We applied tSpace (Fig. 2a) to tonsil and blood B cells, and subsequently used conservative gates based on antibody staining to highlight classically defined B cell subsets for visualization in the tSpace projections (Fig. S4). Cells not falling within the conservative subset gates, which include bridging populations representing transitional phenotypes, are not labeled.

**Figure 2.**
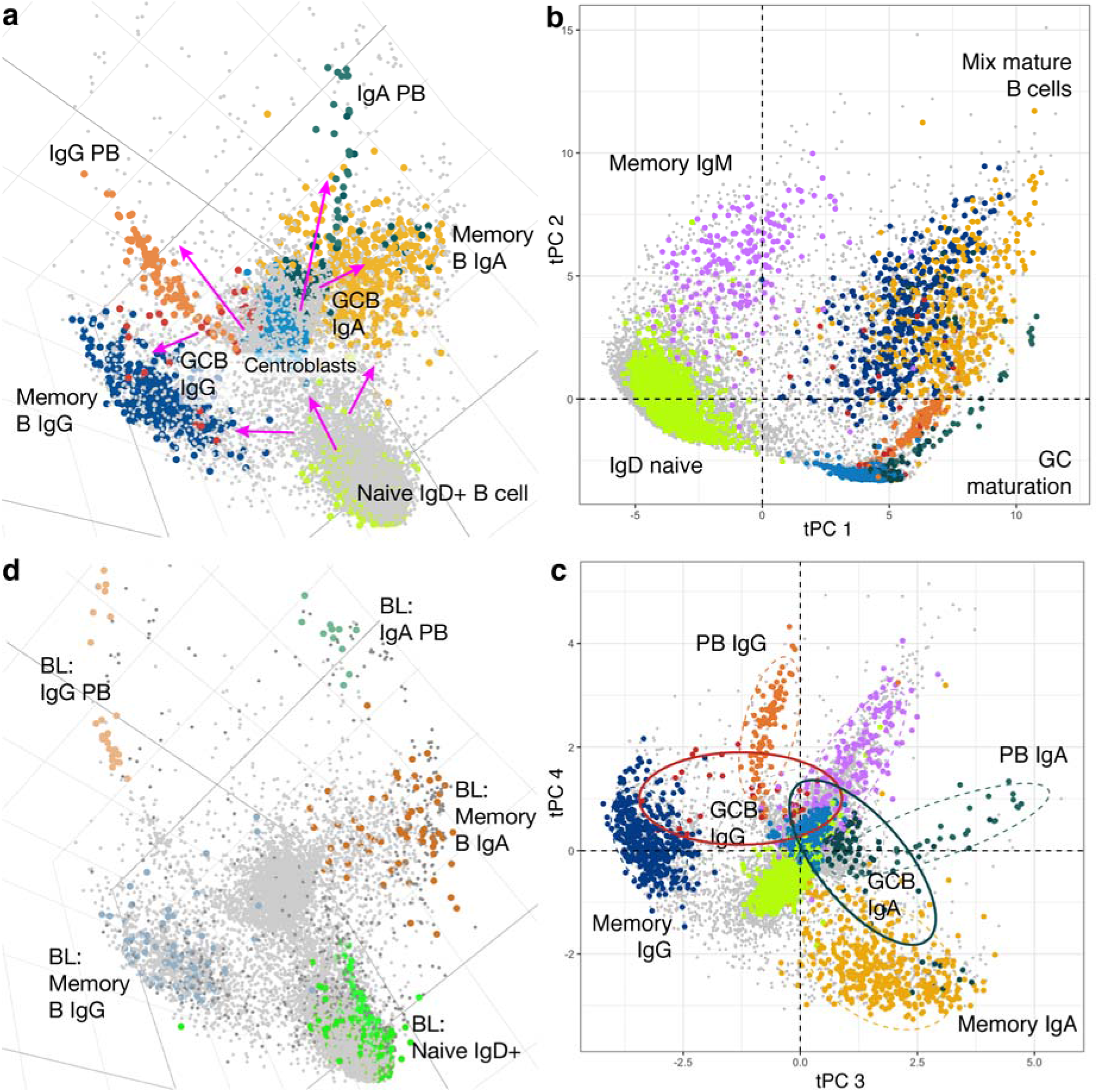
tSpace analysis of B cell differentiation in tonsils and inter-organ trajectories with blood. **a** tSpace unravels maturation paths of B cells starting from naïve B cells in tonsil throughout GC into memory B cells and plasmablasts (PB). Magenta arrows mark suggested directionalities based on known biology. **b-c** Different principal components reveal branches and potential developmental relationships in tonsillar B cell maturation. Ellipses show 80% confidence intervals for indicated clusters. **d** Blood PB align as an extension of tonsillar PB trajectories, while recirculating blood memory B cells overlap with the major tonsil memory cell clouds. In **d**, tonsil B cells are in light gray.

tSpace analysis recapitulates developmental sequences leading from naïve IgD^+^ B cells to tonsil IgG and IgA class switched effector cells. The first trajectory space principal component (tPC1) delineates the transition from naïve to germinal center cells (GCC); tPC2 the differentiation of memory or plasma cells, (Fig. 2b; Fig. S5a); tPC3 and tPC4 pathways to IgA vs IgG class switched cells (Fig. 2c). The distance of cells from naïve B cells within the trajectory space manifold is illustrated in Fig. S5b. A broad strand of cells connects naïve IgD^+^ B cells to proliferating GC centroblasts and centrocytes (Fig. 2a, S5a). Along this path from naïve cells, IgD is downregulated and CD77 is upregulated as cells transition to centroblasts (Fig. S5a, c, d. There are clear, well-delineated trajectories from GCC to class switched PB: CD38, present on activated B cells and GCC, is further induced (Fig. S5c, d) while CD20 is lost (not shown), recapitulating established patterns of antigen regulation in plasma cell development.

The regulation of trafficking receptors during B cell activation, isotype switching and plasma cell generation in human lymphoid tissues has not been resolved. We evaluated chemoattractant and adhesion receptor expression by cells along the trajectory from naïve B cells through the germinal center population to mature plasmablasts. Early naïve cells express CXCR5 which mediates lymphoid follicle homing, CCR6 (Fig. S5c, d) and CCR7 (not shown). These trafficking receptors are downregulated in the transition to germinal center cells, consistent with observations that germinal center cells are non-migratory^7^. CD22 (Siglec2), a B cell specific lectin that moderates B cell activation and also participates in B cell trafficking to gut associated lymphoid tissues^8^, is maintained on GCC but lost during terminal plasmablast differentiation (Fig. S5c, d). Induction of IgA or IgG occurs after initial upregulation of the germinal center marker CD77, consistent with the known role of GC in isotype switching (Fig. S5c, d)^9^. Homing receptors for extra-lymphoid tissue effector sites appear to be induced rapidly upon exit of cells from the germinal center (CD77+) pool. CCR10, a chemoattractant receptor implicated in plasmablast migration to pulmonary and colon mucosae, is upregulated along both IgG and IgA PB lineages, while β7 integrin, a component of the intestinal homing receptor, is highly upregulated in the IgA but not IgG trajectory (Fig. S5c, d). Isotype selective upregulation of tissue specific adhesion receptors within a single inductive tissue has not been observed previously: the mechanisms involved may underlie the selectivity of local IgA secretion for mucosal tissues. CXCR3, implicated in lymphocyte homing to inflamed tissues^10,11^, is coordinately ‘upregulated’ in a minor subset of PB and by memory B cells (Fig. S5a). Many tonsil memory B cells also express CLA, a homing receptor for the vascular addressin CD62E associated with squamous epithelial surfaces including the oral mucosa; CLA was present on some blood plasmablasts (not shown) but was not detected on plasmablast branches in the tonsil.

In contrast to some other tools, tSpace does not constrain or force cells into specific developmental sequences or paths, but instead positions each cell in context with all others even when cell transitions are biologically diffuse. This is illustrated by the dispersed distribution of class switched IgG^+^ and IgA^+^ memory B cells in trajectory space (Fig. 2c, Fig. S5a; and best visualized in 3D embedding – Supplementary Movie 1): memory cells constitute a “cloud” of cells some of which appear to arise from the GC pool as mentioned, while others are closer in trajectory space to the path from naïve B cells to GCs. Cell alignment in trajectory space does not intrinsically provide directional information, thus cells bridging the main memory cell population with germinal centers may reflect recruitment of memory cells into the active germinal center, or generation of memory cells from the germinal center reaction. The surprising alignment of many IgG and IgA expressing cells between naïve and memory populations (adjacent to the naïve to GC path; Fig. S5a) suggests that, in steady state human tonsil, activated B cells may undergo IgA or IgG class switching and conversion to memory cells without transiting through the GC reaction. While class switch recombination is normally attributed to the GC reaction, in some mouse models class switching can occur prior to GC formation, and it is observed in T-independent B cell responses as well^12^. Low expression of CD27 (Fig. S5a) and retention of naïve markers CCR6 and CXCR5 on the class switched cells adjacent to the “naïve to GC” sequence is consistent with this interpretation (not shown). In contrast to their IgG and IgA class-switched counterparts, IgM memory cells (CD27^+^, CD38^−^) are more closely connected to naïve (IgM^+^, IgD^+^, CD27^−^, CD38^−^) cells in most tPCs, with tPC2 specifically expanding this trajectory (Fig. 2b, Supplementary Movie 1). Thus, tSpace recapitulates known pathways of tonsil B cell development and differentiation, presents evidence that human B cells can follow alternative developmental paths that have only been described in animal studies, and reveals developmental stage(s) and transitions at which tissue- and inflammation-specific trafficking receptors are induced.

Developing plasmablasts generated in lymphoid tissues leave their sites of antigen activation and circulate via the blood to distant effector sites. We reasoned that trajectories might link terminally differentiated cells, ready to exit their site of generation, with progeny cells in blood. Indeed, when we applied tSpace to combined blood and tonsil B cells datasets, blood PB aligned at the termini of tonsillar IgG and IgA PB branches (Fig. 2d, Supplementary Movie 2). In contrast to the unidirectional path of maturing plasmablasts, blood memory B cells and naïve IgD^+^ B cells exchange between blood and lymphoid tissues through recirculation. Consistent with intermixing, these subsets overlap extensively with their tonsillar counterparts in trajectory space (Fig. 2d). These results show that tSpace can unfold inter-organ transitions and elucidate developmentally programmed migration patterns of immune cells in settings where experimental analyses of leukocyte trafficking are challenging, as in humans.

Single cell RNAseq is emerging as a powerful tool for the characterization of cell populations and provides rich cellular profiles for studying cell relationships. We applied tSpace to published scRNAseq data from mouse intestinal epithelial cells using 2420 variable genes^13^ (Methods). Intestinal epithelium forms the single-cell layer separating the lumen of small intestine from intestinal lamina propria. Almost all cells in the epithelium have a short life-span of about 4-7 days^14^ and continuous renewal is driven by division of Lgr5^+^ crypt base columnar (CBC) cells residing in the bottom of the intestinal crypts. Transit-amplifying (TA) cells within the crypt differentiate into absorptive (enterocyte) or secretory [goblet, Paneth, tuft and enteroendocrine (EE) cell] lineages. tSpace delineates absorptive/enterocyte and secretory/EE developmental paths (Fig. 3a, Fig. S6a-b). Goblet and Paneth cells define short branches from the proliferating TA pool.

**Figure 3.**
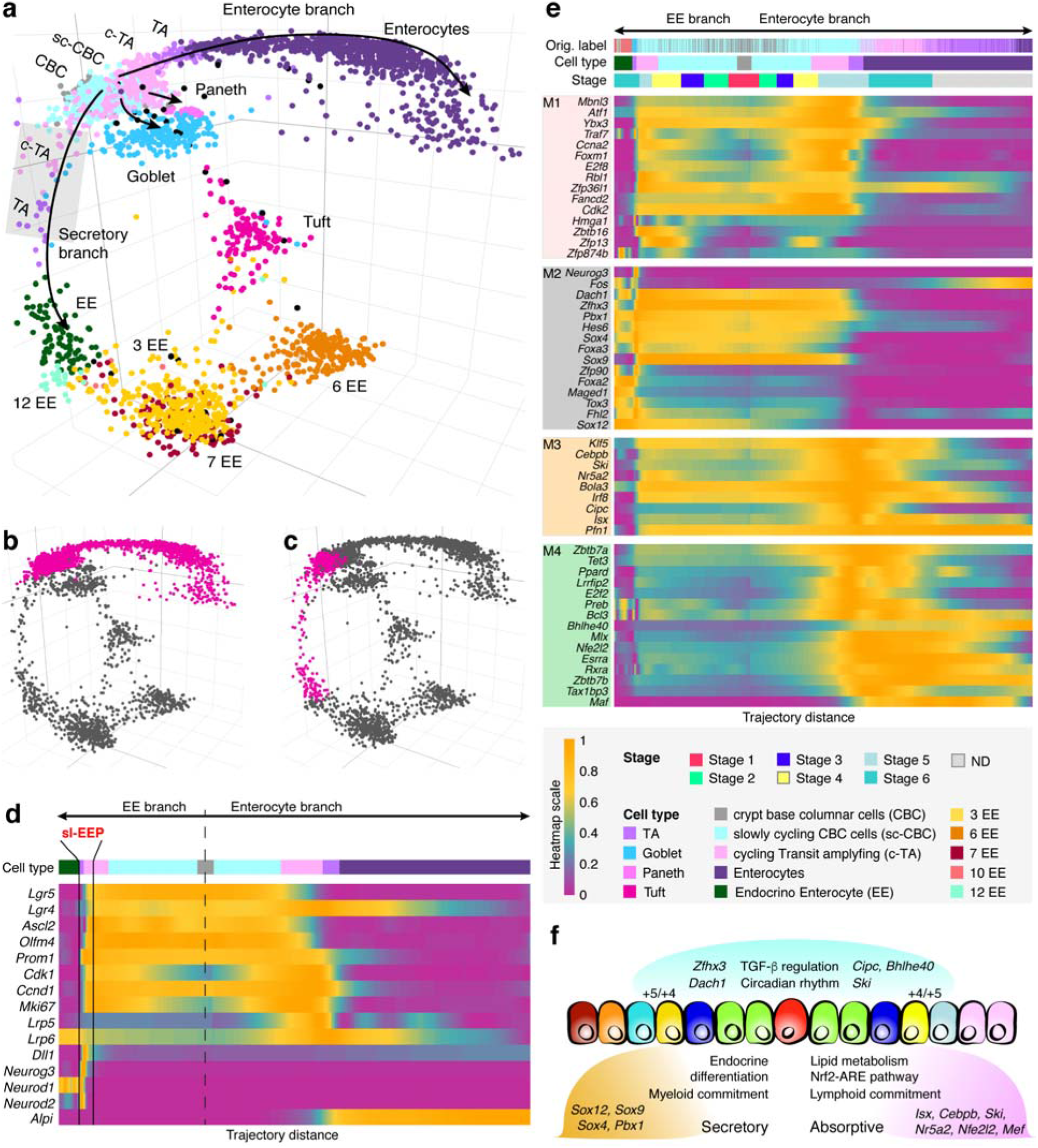
tSpace analysis of mouse small intestinal epithelial cells. **a** tSpace separates trajectories to enterocytes, enteroendocrine (EE), Paneth and goblet cells. CBC and TA subsets were defined by our analysis, as described in Fig. S7., other subsets are labeled as in Yan *et al.*^13^. Shaded rectangle highlights the position of short-lived EE progenitors (slEEP) cells. **b** Isolated enterocyte trajectory. **c** Isolated EE trajectory. **d** Expression patterns of selected genes^15^ (known markers or regulators of intestinal crypt development; expanded gene list in Fig. S8a) along the isolated trajectories. **e** Four detected transcription factor modules in early trajectories, identified by comparing gene expression between cells at similar stages in the two trajectories (Methods): M1 comprises TFs involved in cell cycle and genome integrity expressed in precursor populations (early in the shared trajectory). M2 and M3-M4 differentiate the two lineages and comprise TF’s that may determine cell fate or specialization (see text). Cell stage (Methods) and cell identities defined here (Cell type) or in Yan *et al.* (Original labels) are indicated above the heatmap. ND -fully differentiated enterocytes, not used in trajectory alignment. **f** Summary of differences between two branches suggested by gene regulation along the trajectories. Different genes in the TGF-β and circadian rhythm pathways are expressed in the two lineages (genes in blue above the cartoon). TFs enriched in the EE branch are involved in endocrine secretory cell development; while TFs associated with enterocyte commitment include regulators of lipid/cholesterol metabolism. Expression of *Dll1* and *Sox4* in EE development and *Alpi* in enterocyte differentiation, mark specific progenitor cells located within the +4/+5 position in the intestinal crypt according to the literature^21,26,31^; clear peaks are seen in their expression along the trajectories (Fig. 3d-e) near the TA to differentiated cell transitions, likely representing these specific progenitor populations.

The ability of tSpace to position all cells in developmental relationships allows additional interesting insights. tSpace trajectories reveal a common ‘trunk’ leading to secretory and absorptive branches, but many sc-CBC and c-TA cells (defined Fig. S7) actually segregate to the early EE or enterocyte branches, suggesting that they are already developing towards if not committed to EE or enterocyte fates. To illustrate the application of tSpace to explore developmental progression of gene expression in this context, we isolated trajectories within the tSpace distance matrix starting from the Lgr5^+^ CBC cell population, gated on cells of the enterocyte branch and cells within the early segment of the EE branch preceding EE3 (Fig. 3b-c), and plotted gene expression of cells vs their trajectory distance from CBC cells (Fig. 3d). We focused initially on genes for known hallmarks of intestinal differentiation (Fig. 3d, S8a)^15^. The analysis confirms *Ascl2*^16^, *OlfM4*^17^ and *Prom1*^18^ as robust markers of the presumptive crypt populations (CBC to TA cells) and reveals that the expression of *Prom1* extends into the TA pool, confirming previous findings^19^ (Fig. 3d, S8a). The Wnt agonist *Lgr5* and its homolog *Lgr4* are in resting CBC and dividing slow-cycling CBC (sc-CBC) cells, but the analysis shows that *Lgr4* expression is retained in post-mitotic cells differentiating towards absorptive enterocytes from cycling TA (c-TA), suggesting that in addition to its known role in proliferation of TA cells^20^ it may contribute to enterocyte fate or specification.

Further examination revealed *Dll1*-expressing cells in trajectory space between CBC cells and mature EE populations (Fig. 3a, shaded grey rectangle, Fig. 3d). These cells express genes that define short-lived secretory progenitors (slEEP)^21^, which upregulate EE lineage specification genes *Neurog3*, *Neurod1* and *Neurod2* (Fig. 3d)^22,23^. Consistent with their location in tSpace projection, sIEEP are well-documented precursors of EE cells^14,21,22^. The EE branch proceeds through EE3 cells, recently identified as EE intermediates, giving rise to specialized mature enteroendocrine subsets^13^ (for cell labels see Methods and Fig. S7). Interestingly, sparse intermediates link a single tuft cell population to both CBC/TA and to EE3 cells (best visualized when UMAP is applied to trajectory space matrix, Fig. S6a-b). While the number of intermediate cells linking these two pathways to tuft cells would suggest caution in interpretation, it is noteworthy that a dual origin of tuft cells (directly from Lgr5^+^ CBC cells but also from EE3 enteroendocrine cells) has been proposed from multiple lines of evidence^13,24^. tSpace performed well in comparison with SPADE, a minimum spanning tree (MST) algorithm applied to visualize trajectory relationships in the original analysis of this scRNAseq dataset. SPADE^13^ and tSpace both delineate the major CBC to enterocyte and EE branches, the relationship of goblet and Paneth cells to CBC/TA, and the terminal branching of EE subsets. SPADE generates a 2D representation of an MST structure, an approach that is inherently challenged by non-tree-like developmental paths, such as the paths from EE3 and CBC that converge on tuft cells (Figs. 3a, S6a-b). While tSpace identified a single tuft cell pool with dual connections, SPADE analysis forced tufts cells into two disconnected populations, one arising from CBC cells and the other from intermediates leading to EE3 cells. SPADE also failed to detect or properly position slEEP on the path to EE cells.^13^ In contrast to tSpace which positions each cell in trajectory space, SPADE and related MST algorithms rely on prior definition of cell clusters and limited gene sets, features which run the risk of missing or mislabeling important cell intermediates^13^. sIEEP were defined either as cycling CBC or goblet cells in the published analysis and were subsumed in biologically inappropriate clusters (Figs. S7a, b, d, S8a see original labels).

Overall, cell positioning in trajectory space and the patterns of gene expression reflect observations from decades of research on intestinal development, but also suggest refinements to current understanding. Many TA cells express gene programs leading to secretory vs. absorptive phenotypes, (Fig. 3d, S8a), indicating that fate selection is already initiated within the dividing (TA) pool that arises from Lgr5^+^ CBC. A global survey of transcription factor (TF) expression during early specification of secretory vs absorptive fates has not been described. We evaluated gene expression along tSpace-defined developmental sequences to identify TF that might specify, and/or control downstream cell specialization. Four distinct TF modules were identified (Methods) based on their patterns of regulation along early EE or enterocyte branches (M1-M4, Fig. 3e, S8c). Genes for proliferation and DNA maintenance (M1, e.g. *Ccna2*, *Cdk2*, *Fancd2*, *Rbl1*) are expressed by dividing sc-CBC and TA “early” along the trajectory, as expected. A second module of TF genes is also expressed by early cells but is maintained selectively in the EE branch: these include TF’s associated with endocrine and pancreatic development (e.g. *Foxa2*, *Foxa3*, *Neurog3*, *Sox4*, *Sox9*) that may coordinate secretory pathways within intestinal enteroendocrine cells^25^. Interestingly, among these, tSpace revealed an unexpectedly high and selective expression of *Sox4* in slEEP cells, suggesting it as a novel candidate contributor to EE specification (Fig. 3e, S7c, e): this prediction has been subsequently confirmed^26^. Modules 3 and 4 TFs are expressed preferentially in the enterocyte branch. They include TF involved in lipid and cholesterol metabolism required for mature enterocytes (e.g. *Cebpb*, *Klf5*, *Nr5a2*, Fig. 3e, S8c)^27,28^, but also *Nfe2l2* and *Maf* associated with the activation of *Nfe2l2/Nrf2*-antioxidant response element (ARE) pathway^29^. Enterocytes utilize short fatty acids as a source of energy, and fatty acid metabolism generates reactive oxygen species (ROS), and ROS are also abundant in the intestinal lumen^30^. Upregulation of the *Nfe2l2*-ARE pathway may help protect differentiating enterocytes from oxidative damage^30^. The analysis also identified *Isx* and *Ski* (both within M3) as putative novel markers of TA cells within the early enterocyte developmental branch; lack of specific markers has hindered isolation of TA cells and further probing of their plasticity.

The tSpace approach is conceptually similar to isomap^32^. Both methods provide a global approach to dimensionality reduction, designed to preserve manifold geometry at all scales. Both algorithms determine geodesic distances along a KNN graph. Isomap embeds the resulting distance matrix in low dimensions using multidimensional scaling (MDS). It has been successfully applied to diverse high dimensional datasets^33–36^, but it has not been adopted for high dimensional single cell analyses, perhaps because of well-described limitations. The algorithm is computationally and memory expensive. This has been addressed in part by ‘landmark isomap’ by calculating approximate distances using a set of randomly selected ‘landmark’ cells. To ensure uniform sampling of the manifold, we modify this approach in tSpace by selecting individual cells from each of T K-means clusters, where T is the number of trajectories to be calculated. We show that linear trajectories (distance vectors) calculated from 100 – 250 well-distributed starting cells are sufficient to recapitulate cellular relationships in each of the datasets here. Isomap suffers also from sensitivity to “short circuit” errors if K is too large or if noise in the data positions cells aberrantly between valid branches or populations in the manifold. Short circuits pose a problem with the Dijkstra algorithm, used in isomap to calculate shortest paths between cells. We take advantage of the Wanderlust, which refines distances and avoids “short circuits” by using subgraph averaging and weighting in shortest path calculations based on waypoints^1^. We show that tSpace with Wanderlust improves the definition of developmental paths (Fig. S1). While, isomap uses MDS, a memory intensive algorithm, for dimensionality reduction and visualization of manifold relationships, tSpace utilizes PCA and/or UMAP. Namely, trajectory space matrix is a matrix of the quantitative relative distances from every cell in the data, regardless of its lineage or terminal fates, application of inexpensive PCA on that type of a matrix recreates multi-branching tree of lineages. The first 3 tPCs often embody the most important developmental branches (with simple branching development), but higher tPCs can also reveal critical biological processes. Many methods allow reconstruction of simple developmental branching sequences, but it is becoming increasingly clear that differentiating cells in development and cancer can and often do retain multi-potency even as they mature. This leads to complex higher dimensional ‘lineages’ or developmental pathways that cannot be represented in 2 or even 3D. In this setting, exploration of PCA projections of trajectory space is valuable, as higher tSpace principal components (tPCs) can reveal additional branching pathways and relationships. We illustrate for example the parallel pathways of IgA vs IgG memory and plasma cell development in tonsil B cells, which dominate the 4^th^ tPC. tSpace also implements dimensionality reduction with UMAP^37^. Although UMAP implements a force directed algorithm that obscures developmental distances, it is an excellent tool for reducing dimensionality to 2-3 dimensions for visualization of the tSpace manifold. We show for our scRNAseq example that UMAP embedding of trajectory space reveals developmental branches better than UMAP embedding of the original gene expression matrix (see supplementary text **Comparison of tSpace and other trajectory inference algorithms**).

A number of other methods for trajectory inference have been described and compared^38,39^. We highlight some of the practical features and limitations of tSpace and of other published algorithms designed for multi-branching and complex manifolds (Supplementary Table 3) and examine the performance of several widely used methods in comparison to tSpace in supplementary comments. Few unsupervised methods exist (we tested Monocle2, p-Creode, DPT, PAGA, and because of its speed and usefulness UMAP), and none combine the scalability and quantitative distance metrics of tSpace. For example, tSpace has advantages over algorithms that use memory intensive MST methods to define branch points, as for large datasets these depend on downsampling of cells or calculation of relationships between clusters (rather than individual cells) to reduce computational complexity. Examples include slingshot^40^, p-Creode^41^ and SPADE^42^. As highlighted above in discussion of published analysis of intestinal epithelial cells, downsampling in MST-based methods holds inherent risks of obscuring important cell subsets, and in most algorithms fails to position each cell in developmental relationships. tSpace avoids the loss of individual cell resolution associated with cell downsampling, while retaining the ability to reveal the developmental relationships of all cells to each other. Algorithms that focus on computationally defining branchpoints and tree structures (e.g. Monocle2, slingshot) can also limit appreciation of alternative pathways of differentiation represented by cells that bridge between dominant pathways (e.g. converging paths of tuft cells). Moreover, in contrast to algorithms that rely only on local cell relationships or that use force directed graph methods, the global approach of tSpace estimates distant as well as nearby cell relationships within the manifold. Indeed, the algorithm exports a dense matrix of meaningful cell-to-cell distances that represent measures of the extent of phenotypic change along developmental pathways. As illustrated in our examples, cells along specific developmental pathways and branches can be easily gated (isolated) in plots of tPCs using commonly available software such as Flowjo, JMP or in R (Methods). Trajectories starting from branch termini or other desired points within pathways are readily identified within the tSpace matrix, and plotted vs gene/protein expression to characterize changes in cell phenotypes along isolated developmental sequences (as in Figs. 1–3).

In conclusion, we have presented the concept of trajectory space and its implementation in the tSpace algorithm for elucidation of branching or convergent developmental pathways and mechanisms from single cell profiles. tSpace performs well across different biological systems and platforms and reveals known and novel biology. tSpace embodies a combination of useful features including 1) applicability to any type of data (proteomic, transcriptomic, etc.); 2) simplicity of use; 3) scalability and independence from the need for downsampling; 4) robustness to input parameters; 5) positioning of each individual cell in developmental relationships (allowing visualization of alternative or minor pathways of differentiation); 6) retention of global as well as local cell relationships with export of quantitative measures of cell to cell distances in the manifold; and 7) independence from requirements of clustering or prior information. The tSpace outputs are reproducible, intuitive and amenable to exploration of biology (gene or protein expression, trajectory isolation, etc.). We believe that tSpace will prove useful to the rapidly growing field of singe cell analysis.

## Acknowledgments

Mass cytometry analysis for this project was done on Cyrano instrument in the Stanford Shared FACS Facility, obtained by S10OD016318-01 NIH grant. We thank Raghav and Durga Ganesh for assistance with parts of the code, Menglan Xiang and Sofia Nordling for help with evaluations of the algorithm, Steven Schaffert for constructive discussions and Lourdes Magalhaes for administrative assistance.

