## Supplementary Materials & Methods for "Exploration of cell development pathways through high dimensional single cell analysis in trajectory space"

**Formal description of the tSpace algorithm**

Every analyzed cell c*_i_* is defined by a vector of expression values for each measured marker (v) c*_i_* = (v_1_, v_2_, v_3_, …, v*_j_*), where *i* is between 1 and the total number of cells (N) and *j* is the number of measured markers (m). Profiles of all cells within a sample form an *N-*by-*m* input matrix. Prior to application of tSpace, the input matrix has to be cleaned of outliers and artifacts and adequately transformed, e.g. logicle for FACS data, the inverse hyperbolic sine function for CyTOF and logarithmic for scRNAseq.

In the *trajectory space* concept each cell is represented as a vector of distances to every other cell in the data set. Implementation of the concept through the tSpace algorithm involves three steps.

In the first step tSpace calculates a K-nearest neighbor (KNN) graph, in which there is an edge from cell c_i_ to c_j_ if and only if the distance between c_i_ and c_j_ is one of the K smallest distances between c_i_ and all other cells. The user-defined neighborhood size (K), preferably small, is selected to allow inclusion of every cell in the graph. Values of K ~ 20-30 works for most data sets in our analysis. In order to reduce “short-circuits”, which may occur in basic KNN graph, L (L < K) number of connections for every cell is preserved and a L-KNN sub-graph is created. User defines how many sub-graphs will be calculated by defining a parameter *G*.

The second step of tSpace is computation of a *trajectory space* distance matrix. This step utilizes parallelization for performance improvement. tSpace computes distances [d*_Wanderlust_*(*x*, *y*)] from each cell to every other cell within the L-KNN graph using a modified Dijkstra algorithm, originally published under the name Wanderlust^1^. Briefly, for each cell c*_i_*, Wanderlust initially calculates interim trajectories for each of G sub-graphs. These interim trajectories are averaged to generate a consensus distance to every cell. Cell distances within each sub-graph are calculated in relationship to both the start cell and to “waypoint cells” that are randomly picked from within uniformly distributed subsections of an initial trajectory. The number of waypoints (WP) is user defined. Distances from the start cell to every other cell are iteratively refined using the waypoints and an exponential weighting scheme until convergence (reaching correlation of 0.9999 between the trajectories). The weight matrix is determined using the distances between waypoints and the rest of the cells; this improves calculated distances by giving more importance to the distances around neighboring waypoints. The final *trajectory space* matrix M is a dense matrix of refined distances (d) from each cell to every cell in the dataset, where M*_ij_ =* d*_Wanderlust_*(c_i_, c_j_), i & j ∈ [1, N], with N being the total number of measured cells, and each cell is described by a vector of distances c*_i_* = (d_1_, d_2_, d_3_, …, d*_j_*), i & j ∈ [1, N]. We refer to the vector of distances from cell ci to every other cell as a trajectory.

Lastly, dimensionality reduction (e.g. PCA or UMAP) is used to visualize cells and their relationships in trajectory space.

In the ideal situation, the number of trajectories T is equal to the number of analyzed cells (N). However, when datasets are large calculation of trajectories starting from every cell can be impractical. In this case, trajectories can be calculated starting from a subset of cells that are selected to be representative of all regions and subsets in the population. We find that 100-1000 trajectories are sufficient for accurate determination of branch points and different developmental paths in all samples we have studied to date (Fig. S1). Phenotypically diverse starting cells, representative of all subsets in the population, are identified by selecting one cell from each of a number of T K-means clusters. The number of clusters is equivalent to user defined number of trajectories (T). Thus, the *trajectory space* matrix (M) can be defined as M = (d*_ij_*), i ∈ [1, N], j ∈ [1, T], and each cell is described by a vector of Wanderlust distances c*_i_* = (d_1_, d_2_, d_3_, …, d*_j_*), j ∈ [1, T]. MATLAB and R package implementation of tSpace and accompanying documentation are available online (<https://github.com/hylasD/tSpace> & <https://github.com/hylasD/MATLAB_version_tSpace>). A tutorial on application of tSpace is available online (<http://denisdermadi.com/tspace-trajectory-inference-algorithm>) and as a vignette in R package.

**Robustness and effect of parameters**

To address the robustness of tSpace to parameter selection, we examine (i) the effects of Wanderlust vs Dijkstra trajectory calculation for different T (trajectory number) (Fig. S1); and (ii) the effects of varying K (L is maintained at 0.75K as default in these examples) and distance metric (Fig. S2) on tSpace projections of thymic T cell data. 100 trajectories were sufficient to visualize developmental branching in our thymocyte dataset (Fig. S1a-b, N = ~ 95 000 cells) and scRNAseq (Fig. S1c, N = ~3500 cells); with only minor refinements in cell positioning with 1000 trajectories (Fig. S1).

Furthermore, we compare the effect of the varying K, L and different metrics (Euclidean vs Pearson, Fig. S2). The tSpace output is expected to be influenced by K, the number of neighbors in the KNN graph: Large K, by increasing the number of ‘paths’ between cells, can lead to unwanted connections (short circuits) to developmentally more distant or altogether unrelated cells. Conversely, if K is too small the neighborhood graph will be unconnected. We find that developmental relationships predicted for thymocytes by tSpace are surprisingly robust over a range of K (from 15 to 100 in Fig. S2). At K = 100, however, the positioning of terminal Tregs is distorted, with some Tregs beginning to form a bridge to early developmental stages (Fig. S2, asterisks & insets). Thus, we suggest a default to K ~ 25. The parameter L defines the subset of K connections around each start cell to be used for sub-graphs; thus, L must be < K. The ratio of L to K determines the independence of sub-graphs that are averaged to further reduce the contribution of short circuits. We suggest L/K ~ 0.75, a value that works well in all datasets we have analyzed.

tSpace implements Euclidean, cosine or Pearson correlation metrics to define local distances between cells. Comparison of tSpace using Euclidean and Pearson metrics is illustrated in Fig. S2.

**Materials and methods**

**1. Animals**

C57BL/6J male mice, used for thymic T cell isolation and immunofluorescent microscopy, were bred and maintained in the animal facilities of the Veterans Affairs Palo Alto Health Care System, accredited by the Association for Assessment and Accreditation of Laboratory Animal Care. All animal work was approved by the Institutional Animal Care and Use Committee at the Veterans Affairs Palo Alto Health Care System.

**2. Human tissues**

Heparinized peripheral blood (10 - 40 mL) was obtained via venipuncture and processed using Ficoll density gradient centrifugation (Histopaque-1077, Sigma-Aldrich). The interface containing the peripheral blood mononuclear cells (PBMC) was extracted, washed twice with HBSS without Ca^2+^ and Mg^2+^ (Corning), and cryopreserved in FBS with 10% DMSO (10 - 20 millions/vial, 1mL). Human tonsils were obtained from the Stanford Tissue Bank (IRB protocol number 17204). The tonsils were cut into small pieces and the lymphocytes were released with HBSS containing Ca^2+^ and Mg^2+^ (Corning). The cell suspension was spun at 300 g, 4º C for 5 min; pelleted cells were frozen in FBS with 10% DMSO (10 - 20 millions/vial, 1mL). Samples were stored in liquid nitrogen until use.

**3. Immunostaining for FACS and mass spectrometry**

**Flow cytometry of thymic T cells.** Thymic T cells from two C57BL/6 male mice were immunophenotyped using a panel of antibodies to 13 markers involved in T cell development and maturation. Briefly, thymi were homogenized to obtain a single cell suspension. Cells were blocked with Fc-block (1:100) and rat serum (1:50) for 15 min at room temperature. We stained cellular surface in two steps: first primary antibody cocktail for 40 min, followed by 30 min secondary anti-biotin (streptavidin) stain (Supplementary table 1). We fixed cells in 1% PFA for 5 min and stained intracellular content for Foxp3 in permeabilization buffer (eBiosciences) over night.

On average, 100 000 events were collected using BD LSRFortessa™ (BD Biosciences) cell analyzer with 5 lasers. Some antibodies failed to stain significantly above background: these are not discussed further. Antibodies used for tSpace calculations are labeled in bold.

**Supplementary table 1. FACS antibody panel used for profiling of T cells**

| **Primary Antibody** | **Dilution** | **Clone** | **Channel** | **Source** |
| --- | --- | --- | --- | --- |
| **CD90** | 1:400 | 53-2.1 | APC | BD Biosciences |
| **TCRβ** | 1:200 | H57-597 | AF780 | BioLegend |
| **CD3ε** | 1:200 | 17A2 | BV510 | BD Biosciences |
| **CD4** | 1:200 | RM4-5 | BV711 | BD Biosciences |
| **CD8α** | 1:200 | 53-6.7 | eVolve 655 | eBioscience |
| **CD24** | 1:200 | M1/69 | BUV737 | BD Biosciences |
| **CD25** | 1:200 | PC61 | PerCP-Cy5.5 | BioLegend |
| **CD44** | 1:200 | IM7 | eFluor 450 | eBioscience |
| **CD62L**-biot | 1:200 | MEL-14 |  | BioLegend |
| **CD117** | 1:200 | 2B8 | BUV396 | BD Biosciences |
| **CCR9** | 1:200 | CW-1.2 | FITC | BioLegend |
| NK1.1 | 1:200 | PK136 | PE-Cy7 | BioLegend |
| Streptavidin | 1:200 |  | BV786 | BD Biosciences |
| **Foxp3** | 1:200 | FJK-16s | PE | eBioscience |

**Mass spectrometry (CyTOF) antibodies.** Mass cytometry antibodies were either purchased from Fluidigm (Sunnyvale, CA) or labeled in-house using MaxPAR X8 antibody conjugation kits (Fluidigm) following the manufacturer’s protocol. Mass cytometry antibodies used are in Supplementary table 2. Antibodies were diluted in Candor PBS Antibody Stabilization solution (Candor Bioscience GmbH, Wangen, Germany) supplemented with 0.02% NaN_3_ (final concentration) and stored up to 6 months at 4º C.

**Mass cytometry of human B cells in tonsils and blood.** Cryopreserved PBMC or lymphocytes from tonsil were thawed at 37º C with gentle manual agitation, washed twice with HBSS without Ca^2+^ and Mg^2+^ supplemented with 10% bovine calf serum and resuspended at 20M cells/ml in CyFACS buffer (PBS with no heavy metal contaminants (Rockland Immunochemicals, Limerick, PA) with 1% BSA (Sigma) and 0.1% sodium azide). 5 million cells of each sample were used for staining. Cells were initially blocked for 10 minutes at 4º C with 0.5 μl normal human serum (Sigma) and 2.5 μl goat serum (Gibco) and then washed in CyFACS buffer (all subsequent washes were done with CyFACS buffer unless specified otherwise). Cells were incubated with purified, unconjugated anti-GPR15 for 45 minutes 4º C, washed once, then incubated with goat-anti-mouse IgG -168 for 30 minutes 4º C and washed. Cells were blocked with 2.5 μl normal mouse serum (Sigma) for 10 minutes 4º C, washed once and stained with a cocktail of primary antibodies for 45 minutes 4º C. Samples were washed and stained with metal-conjugated secondary antibodies against FITC to detect CLA-FITC and APC to detect α4β7-APC for 30 minutes 4º C and washed. Live/dead stain was performed for 30 minutes 4º C using139In-DOTA maleimide (Microcyclics, Plano, TX) diluted 1:2000 in CyPBS (Rockland), washed twice and fixed overnight at 4º C in 2% paraformaldehyde (Electron Microscopy Systems). The next day, samples were washed twice and barcoded using the 20-plex Cell ID kit from Fluidigm according to the manufacturer’s protocol except for two modifications. Cells were fixed overnight in 2% PFA instead of fixed in FixI buffer (for 10 mins at RT) provided with the kit and barcoding was performed after surface staining rather than before. After barcoding, the samples were washed once in CyFACS, once in fixation/permeabilization buffer (eBiosciences) and then combined prior to intracellular staining. Intracellular staining was performed for 45 minutes 4º C using a cocktail of antibody conjugates against intracellular targets, washed twice with CyFACS buffer and DNA content stained for 20 minutes at room temperatureusing Ir-interchelator (Fluidigm) diluted 1:500 in 2% PFA. Final washes were performed (twice with CyFACS, once with CyPBS and once with milliQ water) then resuspended in 500 μl milliQ water.

The samples were filtered through cell strainer cap FACS tubes, counted and the cell concentration adjusted to 600 000 - 1 million cells/ml with milliQ H_2_O containing bead standards (Fluidigm) prior to acquiring the sample on the CyTOF 2.0 mass cytometer (Fluidigm). Between 1 - 1.5 million total events were acquired. After the samples were manually de-barcoded by gating in FlowJo using the specific combination of three metals unique to each individual sample, 30 000 - 70 000 total cellular events remained.

**Supplementary table 2. Mass cytometry antibodies**

| **Mass** | **Primary Antibody** | **Conc. (ug/million cells)^1^** | **Clone** | **Source** |
| --- | --- | --- | --- | --- |
| 141-Pr | **CCR6** | 0.5 μl | G034E3 | Fluidigm |
| 142-Nd | **CD19** | 0.13 μl | HIB19 | Fluidigm |
| 143-Nd | **IgD** | 0.0035 μg | IA6-2 | Biolegend |
| 144-Nd | **CXCR5** | 0.02 μg | RF8B2 | Biolegend |
| 145-Nd | CD4 | 0.12 μl | RPA-T4 | Fluidigm |
| 146-Nd | CD8 | 0.01 μg | SK1 | Biolegend |
| 147-Sm | **CD20** | 0.13 μl | 2H7 | Fluidigm |
| 148-Nd | **CD69** | 0.15 μg | FN50 | Biolegend |
| 149-Sm | **CCR4** | 0.08 μl | 205410 | Fluidigm |
| 150-Nd | CD3 | 0.01 μg | UCHT1 | Biolegend |
| 151-Eu | **CD103** | 0.05 μg | BerAct8 | Biolegend |
| 152-Sm | TCRγδ | 0.13 μl | 11F2 | Fluidigm |
| 153-Eu | **CLA** | 0.04 μg | HECA452 | Biolegend |
| 154-Sm | **CCR2** | 0.05 μg | K036C2 | Biolegend |
| 155-Gd | **CD22** | 0.05 μg | HIB22 | Biolegend |
| 156-Gd | **CXCR3** | 0.13 μl | G02SH7 | Fluidigm |
| 158-Gd | CD14 | 0.04 μg | M5E2 | Biolegend |
|  | CD33 | 0.04 μl | WM53 | Fluidigm |
| 159-Tb | **CD77** | 0.09 μg | 5B5 | Biolegend |
| 160-Dy | **CCR7** | 0.11 μg | G043H7 | Biolegend |
| 162-Dy | Ki67 | 0.5 μl | B56 | Fluidigm |
| 164-Dy | **CCR9** | 0.2 μg | L053E8 | Produced from Hybridoma |
| 165-Ho | **CD62L** | 0.03 μg | Dreg200 | Produced from Hybridoma |
| 166-Er | **CCR10**^2^ | 0.1 μg | 1B5 | Produced from Hybridoma |
| 167-Er | **CD27** | 0.06 μl | L128 | Fluidigm |
| 168-Er | **GPR15** | 0.33 μg | 367902 | R&D Systems |
|  | goat-anti-mouse IgG | 0.5 μg | 10535 | Invitrogen |
| 169-Tm | **α4β7**-APC | 0.22 μg | Act-1 | Produced from Hybridoma |
|  | Anti-APC | 0.2 μg | APC003 | Biolegend |
| 170-Er | **IgA** | 0.005 μg | Polyclonal | BD Biosciences |
| 171-Yb | **β7** | 0.023 μg | FIB504 | Biolegend |
| 172-Yb | **IgM** | 0.2 μl | MHM-88 | Fluidigm |
| 173-Yb | **P-selectin** **Ig** | 0.13 μg | 137-PS | R&D Systems |
| 174-Yb | **CD38** | 0.06 μg | HIT2 | Biolegend |
| 175-Lu | CD56 | 0.08 μg | HCD56 | Biolegend |
| 176-Yb | **IgG** | 0.03 μg | Polyclonal | BD Biosciences |
| 206-Bi | CD16 | 0.2 μl | 3G8 | Fuidigm |

^1^ amounts used of conjugates purchased from Fluidigm are listed in volume measurements since the concentration is unknown.

^2^ a kind gift from Dulce Soler

Cell markers used for tSpace calculation are highlighted in bold.

**4. Single cell RNAseq of mouse intestine**

In order to test tSpace performance with scRNAseq, we used a previously published dataset of intestinal cell populations^2^. The authors kindly provided normalized and scaled expression matrices for their selected variable genes on our request.

**5. Computational processing of acquired data**

**Software.** tSpace was performed in R [3.4.0 (2017-04-21)] and MATLAB [9.3.0.713579 (R2017b)]. R language was used for other analyses: plots (plotly & shiny packages) and t-SNE (Rtsne package), developmental branches isolation. All analysis code is available upon request. For gating of cell populations, we used FlowJo (10.2).

**Processing and tSpace analysis of flow cytometry data.** We inspected raw FCS files for artifacts and outliers in R or FlowJo and filtered out all events with fluorescence values higher or lower than 0.0001% quantile of the measured markers. We used t-SNE to filter out cells that were negative for all markers to be used in tSpace.

**For thymus**, NK1.1^+^ cells were removed manually in FlowJo. Remaining events were exported in a CSV file for tSpace. Thymic data were transformed using logicle^3^ (included in the MATLAB tSpace package). tSpace parameters were: distance metric = cosine, K = 15, L = 12, G = 5, WP = 20 and T = 1000. K-means clustering, as part of tSpace, on expression values of measured markers was used to define 1000 clusters for selection of the T (1000) trajectory start cells.

**Processing of mass cytometry (CyTOF) FCS files.** Data were normalized using the normalization software embedded within the CyTOF software. CyTOF records a value of zero for channels without detected metal. For better visualization, all zero values were assigned random values from a uniform distribution with minimum -1 and maximum 0 using a script in R. Data was transformed applying inverse hyperbolic sine (asinh function in R and MATLAB) with coefficient 5. We used t-SNE to find outliers, which were gated out in FlowJo. For tonsil, CD19^+^ or CD38^high^ cells were gated for tSpace. Parameters in tSpace analysis were: distance metric = cosine, K = 17, L = 12, G = 5, WP = 20 and T = 1000 (Fig. 2, Fig. S6) or T = 100 Fig. S5.

**Processing of scRNAseq files.** Variable genes (2420) determined by the authors using Seurat R package^4^ were used for tSpace calculation. Parameters used for tSpace were distance metric = Pearson correlation, K = 20, L = 15, G = 5, WP = 15. We calculated ground truth trajectory matrix (for all cells, T = 3521).

**tSpace running times.** We measured the (reasonable) time required for tSpace analysis (Supplementary table 4.) for 100 and 1000 trajectories (T) for our B cell tonsil dataset (17,956 cells, 26 variables), for the 3521 cell/2420 variable gene, 3521 cell/ 40 principal components scRNAseq sample and for our thymus dataset (~95,000 cells, 12 variables). Run times were obtained on a 2-core personal computer. tSpace runs on personal computers and can handle large datasets because it is light on memory and the process is parallelized. The number of trajectories used, cells analyzed, and measured protein or gene parameters all impact computation time, as does the hardware. We show that only 100 - <1000 trajectories are required for analysis (Fig. S1). For cytometry datasets, we generally run tSpace on all measured parameters. For scRNAseq studies, we often run the algorithm with up to 1500 – 2000 genes, or significant principal components. For any larger data, especially in number of cells, we suggest use of multicore computer to speed up analysis.

**Supplementary table 4. Running times of tSpace in minutes for different type of data sets in number of cells, variables and trajectories.**

| **Data set** | **Cell no.** | **Variable no.^1^** | **Trajectory no.** | **Waypoint no.** | **Time [min]** | |
| --- | --- | --- | --- | --- | --- | --- |
|  |  |  |  |  | **R** | **MATLAB** |
| B cell | 17 956 | 26 | 100 | 20 | 40 | 19 |
| B cell | 17 956 | 26 | 1000 | 20 | 289 | 198 |
| intestine | 3521 | 2419 | 100 | 20 | 90 | 52 |
| intestine | 3521 | 2419 | 1000 | 20 | 927 | 393 |
| intestine | 3521 | 40 | 100 | 20 | 6 | 2 |
| intestine | 3521 | 40 | 1000 | 20 | 324 | 23 |
| T cell | 94 807 | 12 | 100 | 20 | 313 | 85 |
| T cell | 94 807 | 12 | 1000 | 20 | 1621 | 860 |

^1^ variables were measured phenotypic markers (FACS/mass spectrometry) for B and T cell data, variable genes and the first 40 principal components for small intestine scRNAseq

**Labeling of literature defined cell populations for tSpace validation.** We manually gated on the indicated populations of T and B cells in FlowJo as illustrated in Fig. S3. and Fig. S4. Gated subsets were used to label and validate cell positions in tSpace visualization. For intestinal data we used cluster annotations from the original study for fully differentiated EE and enterocyte populations, however crypt associated (stem and transit amplifying) populations were labeled based on accepted markers along the developmental trajectories^5^ shown in Fig. S7. We used gene expression along the isolated developmental branches to identify and define crypt base columnar (CBC) cells, slowly cycling sc-CBC, cycling transit amplifying (c-TA) and TA cells. Examination of markers proposed to mark so-called Potten’s “+4 cells”^6^, suggested that these cells likely co-exist with or are the same as sc-CBC cells (Fig. S7c,e). Interestingly, many cells expressing transit amplifying markers and proliferation genes were assigned to already differentiated cells or cycling stem cells in the original publication (Fig. S7a-c). With tSpace we were able to detect, and position in putative developmental sequence, rare and transient populations (e.g. short lived enteroendocrine progenitors, slEEP, Fig. 3d).

**Isolation of developmental branches and calculation of expression changes along them**

We manually gated on cells along specific developmental branches or pathways as visualized in the first 3-5 tPCs, using standard gating approaches in Flowjo, R or JMP. An example can be seen in vignette file accompanying our R package or online (<http://denisdermadi.com/tspace-trajectory-inference-algorithm>). Trajectories are directionless, thus, for orientation we relied on prior knowledge in the context of manually gated cell populations (e.g. thymic T and tonsil B cell developmental sequences) or expression of known marker of stem cells in intestine (*Lgr5*). Trajectory distances along the isolated developmental sequences were accessed from the tSpace distance matrices: we identified trajectories within the tSpace cell distance matrix that ‘start’ from cells at or near the putative origin of isolated sequences (i.e., trajectories for which the chosen start cells have the lowest distance value). Column names of trajectory space matrix contain cell indices associated with each trajectory in the distance matrix, facilitating identification of desired trajectories (see example online: <http://denisdermadi.com/tspace-trajectory-inference-algorithm>). One or more such trajectories were averaged for each developmental pathway isolated.

Cells within isolated trajectories were ordered based on their trajectory distances and smoothed values of gene/protein expression along trajectories were visualized.

**6. Systems biology analysis downstream of trajectory inference**

**Alignment of the intestinal trajectories and transcription factor analysis**

Absorptive and secretory branches share many early cells e.g. CBC, sc-CBC and c-TA (Fig. 3a-c) before they separate fully in tSpace projection. In order to compare transcription factors expressed during initial branching of secretory and absorptive cells, early segments of secretory and absorptive trajectories were aligned using dynamic time warping (DTW)^7^. DTW aligns sets of data points that can be represented as linear sequences, here cells ordered by tSpace along the absorptive and secretory branches. It calculates an optimal match between two sequences independently of rates of change. We performed DTW as a downstream analysis in R using dtw package and all variable genes to align the two branches, with absorptive as template and secretory as a query (Fig. S8b). DTW allowed us to split aligned sequences into 6 stages based on commonalities in gene expression. Significant differences in expression of mouse transcription factors (TFs, <http://tcofdb.org/>) within the same stages between the two trajectories were determined using a permutation test^8^. TF modules were determined based on correlation of expression of significantly different TFs along the aligned trajectories.

**Gene ontology of transcription factors**

Ontology terms of TFs were accessed using the R package Biomart and manually curated for biologically relevant terms highlighted in the Fig. 3. Many of the EE or enterocyte lineage TFs lack significant ontology annotation^9^.

**Comparison of tSpace and other trajectory inference algorithms**

We compared the output of tSpace with p-Creode, Monocle2, diffusion maps, PAGA, and UMAP (Fig. S9-S13), all trajectory inference algorithms designed to be agnostic towards different single cell technologies, and not require any additional input or supervision from the user (e.g. estimates of the relative rates of proliferation and loss, start or end cell points). Each of the algorithms provide different outputs, therefore it is hard to have standardized mathematical comparison, however we use known biology to determine (i) how faithfully each one of them reconstructs developmental relations, (ii) how deterministic output is and (iii) sensitivity of the algorithm to transient and rare populations.

**p-Creode** was originally developed for mass cytometry therefore we hoped to find it easy to apply to our FACS and mass cytometry datasets. We found p-Creode, despite the existence of the available tutorial, difficult to optimize due to striking changes if parameters are slightly altered as illustrated in Fig. S9 (T cell) & S10 (B cell). Moreover, in order to run p-Creode, both of our data sets (94,807 T cells, or 17,916 B cells) had to be downsampled below 14 000 cells, impacting detection of rare cell populations (e.g. in Fig. S9 Foxp3+ Tregs are not separated as a branch of CD4 T cells, and in Fig. S12 plasmablasts are never detected). The inconsistent output of cell cluster relations and marker intensities produced by p-Creode, in our opinion, would significantly hinder interpretation of novel datasets. These observed problems are in fact consistent with the original p-Creode publication, where the authors show the algorithm yields quite different outputs when data is re-sampled^3^.

In contrast, tSpace is highly reproducible and robust. Run on the same sample and with the same parameters, it is nearly deterministic: the only non-deterministic component of the algorithm is the selection of waypoints. It is also robust to the selection of tSpace parameters (Figs. S1 & S2). It is significant that Herring *et al.* independently applied p-Creode to a thymocyte flow cytometry dataset and (just as in our own test) p-Creode failed to define Treg branching. Again, this likely reflects the unavoidable problems associated with the requirement for clustering or downsampling.

**Monocle2** failed to order T and B cells in a developmentally meaningful way (Fig. S11a, Fig. S12a). In addition, Monocle2 also required downsampling of both datasets, probably due to memory reasons.

**Diffusion Maps** ordered cells comparable to tSpace providing developmentally correct relations (Fig. S11b, Fig. S12b). However, diffusion maps do not reveal multiple branches as clearly as tSpace (please compare Fig. 1b and Fig. S11b). Also, in the diffusion map projection IgM memory B cells are positioned as an intermediate state from naïve B cells to memory IgA and IgG B cells, instead of stemming as a separate branch from naïve B cells (the latter expected from known biology). The limited ability to reveal multiple branches is consistent with the original report^4^, in which the authors claim that for multiple subsequent branches, diffusion maps should be applied iteratively. Interestingly, the first diffusion component, for both data sets (mouse T and human B cell), did not reflect developmental relations (Fig. S11c & Fig. S12c).

**Partition-based graph abstraction** (PAGA) relies on existing algorithms to unveil data manifolds, such as t-SNE or UMAP, (Fig. S11d & Fig. S12d) and the Louvain algorithm to determine clusters (Fig. S11d-f & Fig. S12e-f). The “novel step” in PAGA analyses determines meaningful connections between the clusters and partitions the graph. PAGA analysis of T and B cells is summarized in Fig. S11d-h and S12d-g, respectively. While automatic clustering for the most part agrees with manual gating (Fig. S11e-f & Fig. S12f), it fails to subset DP TCRβ+ CD3ε+ (DP*), SP CD8 recent emigrants in T cell dataset, and is not always concordant with expert manual gating, which remains the gold standard in immunology. PAGA analysis of T cells (Fig. S11g), even after optimization, does not connect DN2 and DN3 (cluster 10) to the rest of the T cells, and connects SP CD4 (cluster 7) and SP CD8 (cluster 9) via Treg (cluster 11). Treg should branch from CD4 cells. [Interestingly, at very high K of 100, tSpace begins to make this aberrant connection via Treg as well (Fig. S2; this is discussed as an issue with tSpace parameter selection in the Supplementary Methods).

Increasing the number of clusters in PAGA T cell analysis only fractionates DP T cells and does not improve connectivity between cell populations (Fig. S11h). PAGA analysis of B cells (Fig. S12g) connects clusters similarly to diffusion maps: blood naïve B cells start in cluster 4 and differentiate via clusters 12, 3, 0, 6, 1 into memory IgM cells (cluster 13), or IgG memory (cluster 7), and IgA memory (cluster 9). Memory IgM cells are not connected to germinal center (clusters 5, 14 and 2). Cluster 15, mix of IgA, IgG memory cells connects to cluster 2, which is a mix of IgA GCB, and to lesser extent centroblasts and GCB IgG B cells, already suggesting discrepancy to known biology in which GCB IgA or IgG cells should be differentiated out of centroblasts (clusters 5 & 14). Overall, PAGA does not agree with known biology completely, which can be problematic. Additionally, as much as the abstraction step in PAGA can be useful as a summary, it can also reduce a biologist’s ability to explore the data and isolate fine structures or features seen only in single cell/single point representations. This is amply illustrated in the confusing connectivity of abstracted relationships by PAGA in our tonsil dataset. In contrast, the tSpace projection allows visualization of predominant paths and branches but retains positioning of individual cell intermediates that rather intuitively illustrate alternative paths suggested by the data.

**Uniform Manifold Approximation and Projection** (UMAP) of T cell (thymus) and B cell (tonsil) data unveils the manifold (Fig. S11d & Fig. S12d) and aligns cells in developmentally meaningful ways, similarly to tSpace. However, tSpace analysis of intestinal single cell RNAseq outperforms UMAP in preservation of cellular developmental relations (UMAP clusters cells appropriately but does not retain the lineage relationships. Fig S13a & b).

Although UMAP applied conventionally to phenotypic profiles does not perform as well as tSpace, UMAP offers an interesting tool for visualization of tSpace relationships. Please see and compare UMAP embedding of cells based on tSpace trajectories (Fig. S6 and S13c) and contrast it with UMAP embedding of phenotypic profiles (Fig. S13a & b). UMAP does a good job of reducing trajectory space to two dimensions while preserving relationships and branching.

We also analyzed the intestinal scRNAseq data with diffusion maps, Monocle2 and UMAP (Fig. S13). Overall, the diffusion map algorithm performs well in determining major branches, but fails to unveil early differentiation steps or sub-branching within the absorptive and secretory branches. Additionally, rare populations e.g. slEEP cells (express *Neurog3*), are not easily detectable (Fig. S13d-e). UMAP, depending on the minimum distance parameter (minimum value ~0, maximum value 1) will either show separated clusters (Fig. S13a, min_dis = 0.1) or gradual connections between them (Fig. S13b, min_dis. = 1). Contrary to diffusion maps, UMAP separates rare slEEP cells from goblet cells (Fig. S13b, arrow). However, even with the maximum value for minimum distance parameter, UMAP embedding of cell profiles is not able to show the known connection between slEEP cells and rest of the enteroendocrine subsets.

Taken altogether, tSpace visually represents all intestinal developmental stages more equally than diffusion maps and detects multiple branches better than diffusion maps or UMAP alone. Existing algorithms by virtue of clustering (or use of force directed graphing) also limit the ability to visualize the existence of individual or scattered cells between major paths, which may reflect the diversity of differentiation sequences possible within a population. We believe a major reason for the strong performance of tSpace is the Wanderlust algorithm for calculation of distances, which distinguishes it from all the other methods here. The representation of cells in the dense matrix of trajectory space rather than the sparse matrix of KNN graph space also adds to the robustness of the method.

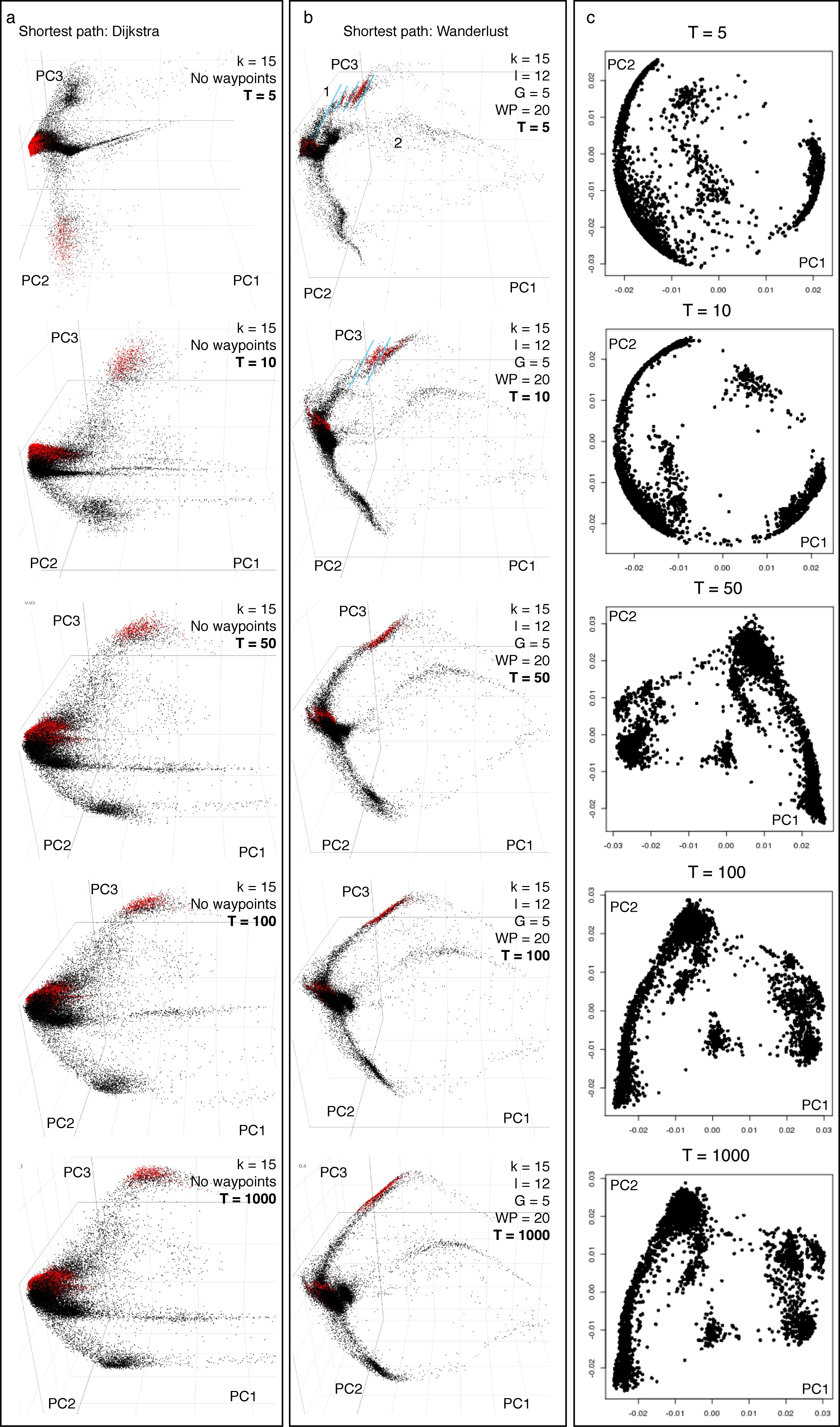

**Fig. S1. Effect of Wanderlust and of number of calculated trajectories (T) on tSpace output: a-b** thymic T cell data and **c** scRNAseq of mouse intestine. **a** tSpace analysis using the Dijkstra algorithm to define distances^10^. The Dijkstra algorithm calculates shortest distances between cells within a KNN, but without the subgraph averaging or waypoint optimization implemented in Wanderlust. An increase in calculated trajectories slightly improves the shape of data. Mathematically, tSpace with Dijkstra is similar to the isomap^11^. **b** tSpace analysis of thymocyte FACS data using Wanderlust^1^. Use of Wanderlust to define trajectories results in tighter and more distinct branches (compare right to left panels). Increasing the number of calculated trajectories improves the definition of developmental sequences as well; but the position of cells in trajectory space (in the tSpace output) tends to stabilize between 100 and 1000 trajectories in experimental datasets. Breaks between cells (blue lines) seen with very low T (5 or 10 calculated trajectories) reflect the influence of waypoints. For orientation, we labeled in red DN3 and DP T-cells. **c** tSpace on intestinal scRNAseq dataset illustrating stabilization of developmental relationships between 100 and 1000 T. For visualization of cell relationships in trajectory space, PCA embedding is used here.

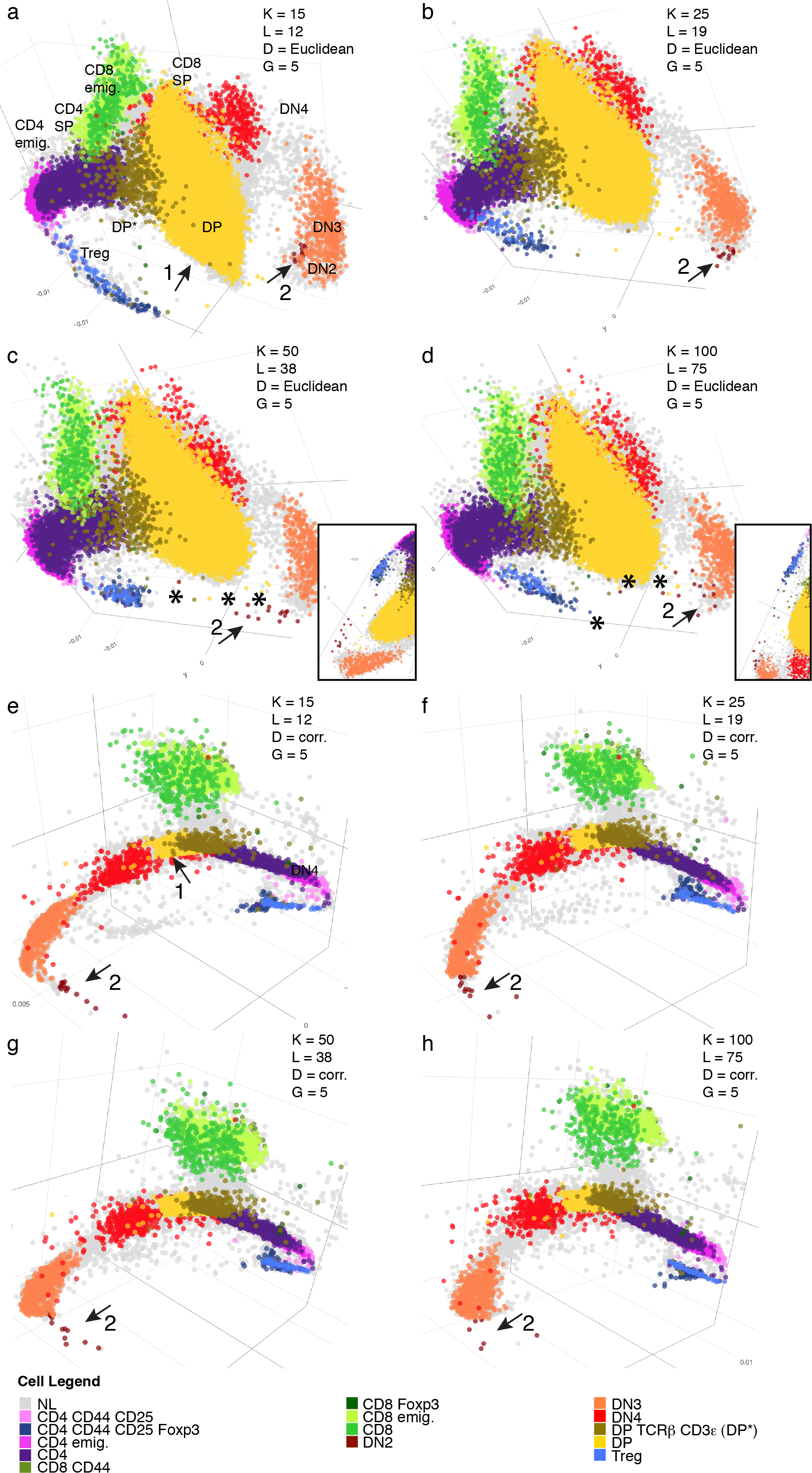
**Fig. S2. tSpace output as a function of *K,* *L,* and distance metric (Euclidean vs. Pearson correlation) a-d** PCA embedding of trajectory space matrices with different parameters using Euclidean distance. **e-f** PCA embedding of trajectory space matrices with different parameters using Pearson correlation. Pearson correlation tightens populations and branches compared to Euclidean (arrow 1). Overall results are robust to a range of parameters, but smaller K reveals more local details and structures of the manifold, and prevents dispersion of cells (e.g. see small DN2 subset, arrow 2). With high K (Euclidean metric), some DN2 cells begin to form a bridge inappropriately towards terminally differentiated Tregs (asterisks and inset panels showing different angle). The choice of distance metric (Euclidean vs Pearson correlation) affects the shape of the manifold but developmental relationships are retained. The numbers of trajectories (T = 250) and subgraphs (G = 5) were constant. Results with 15 subgraphs were similar (not shown).

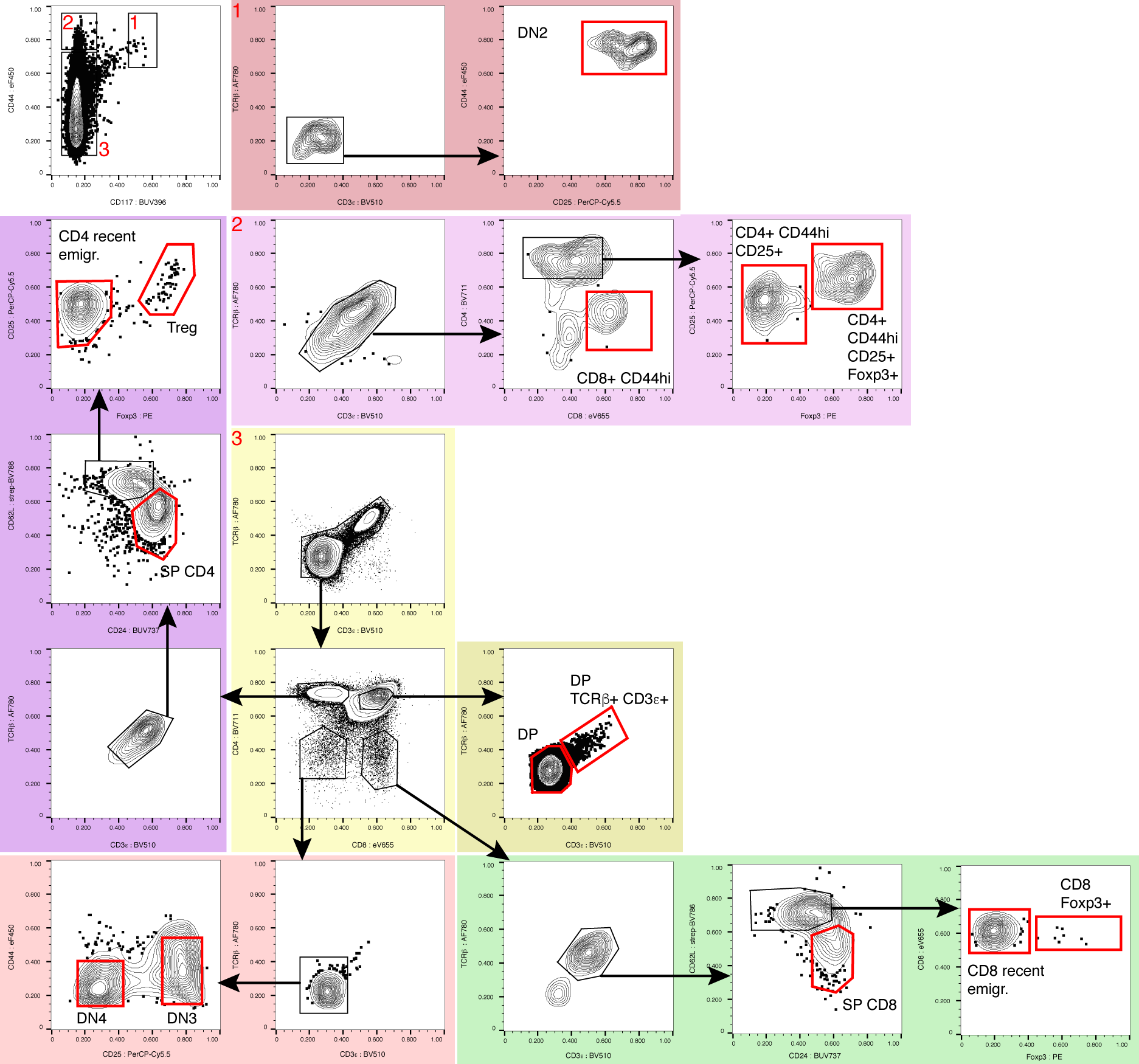

**Fig. S3. Manual gating strategy for conventional definition of T cell subsets in the thymus.** Red gates mark final cell populations used for highlighting in tSpace visualization. Arrows mark flow of the gating.

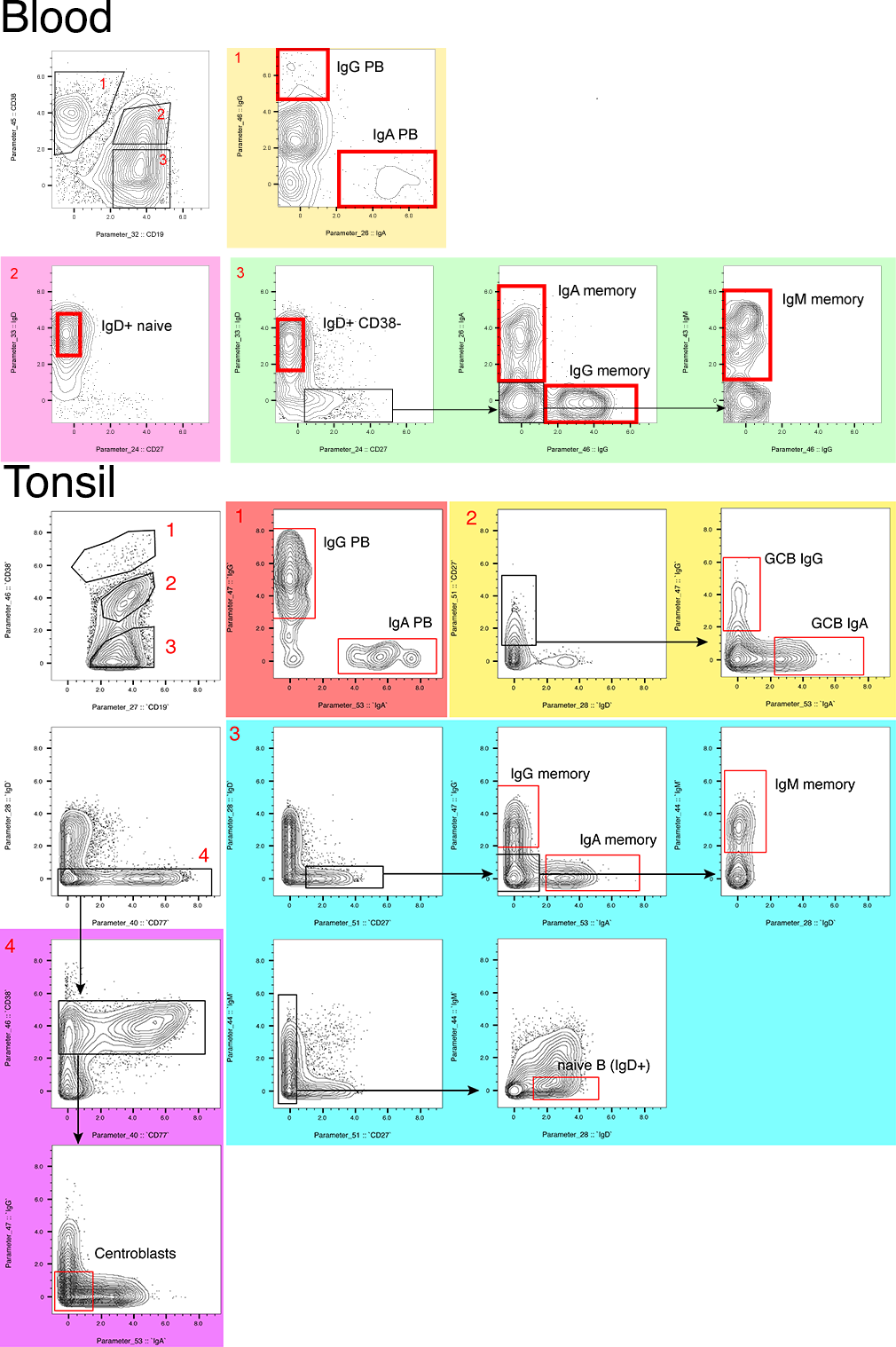

**Fig. S4. Manual gating strategy for tonsil B cell subsets.** Red gates mark final cell populations labeled for visualization in tSpace analyses. Arrows mark flow of the gating.

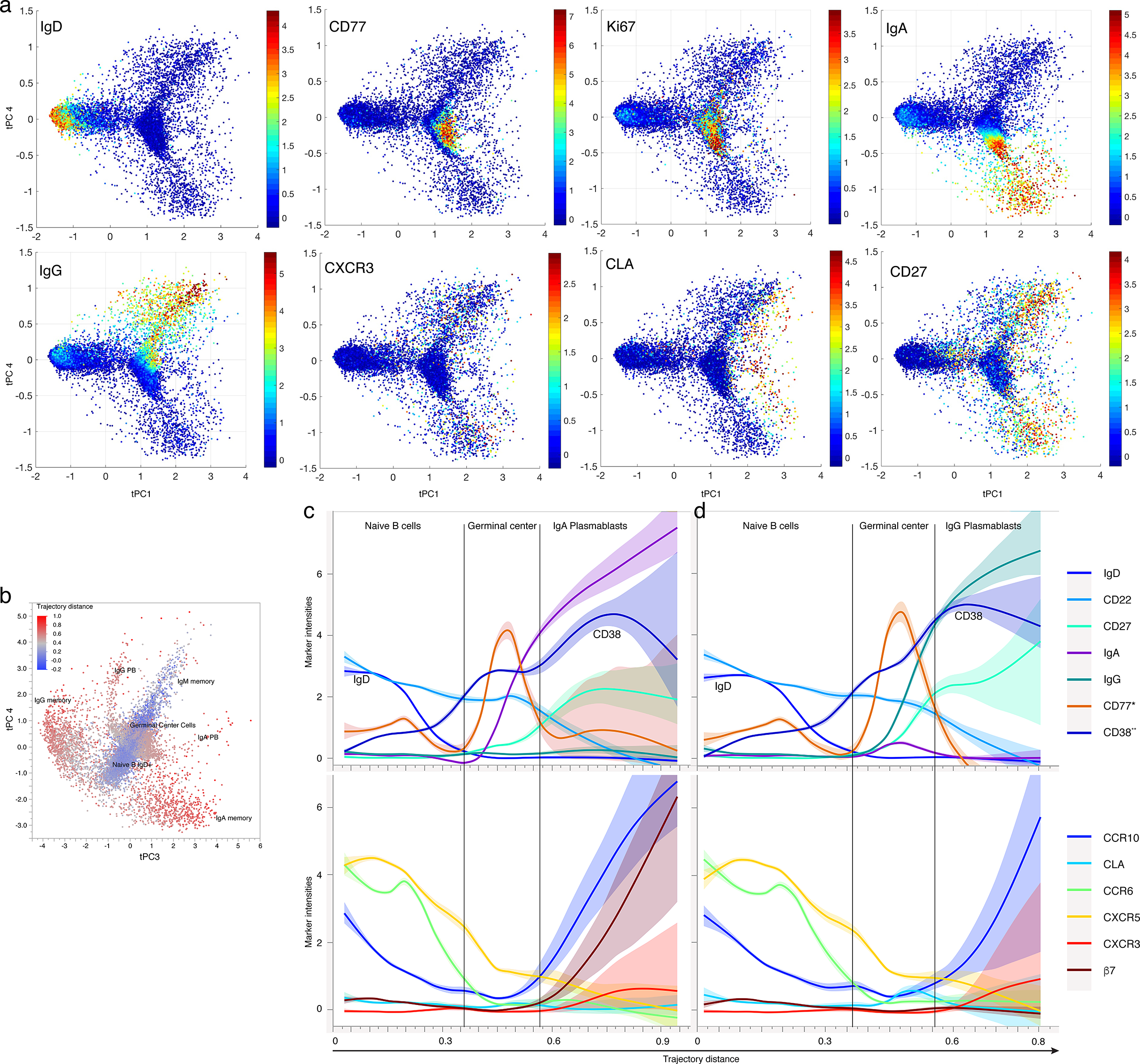

**Fig. S5. tSpace reveals tonsil B cell development. a** Principal component embedding of cells in trajectory space: principal components tPC1 (delineating naïve to GC and mature memory transition) vs tPC4 (delineating pathways to IgG vs IgA expression) are plotted**.** Coloring based on cell staining for the indicated markers. **b** tSpace projection using tPC3 and tPC4 with cells colored by distance from naïve B cells within trajectory space. **c, d** Changes in phenotypic markers and trafficking markers along B cell maturation trajectory from naïve B cells to IgA (c) or IgG (d) plasmablasts.

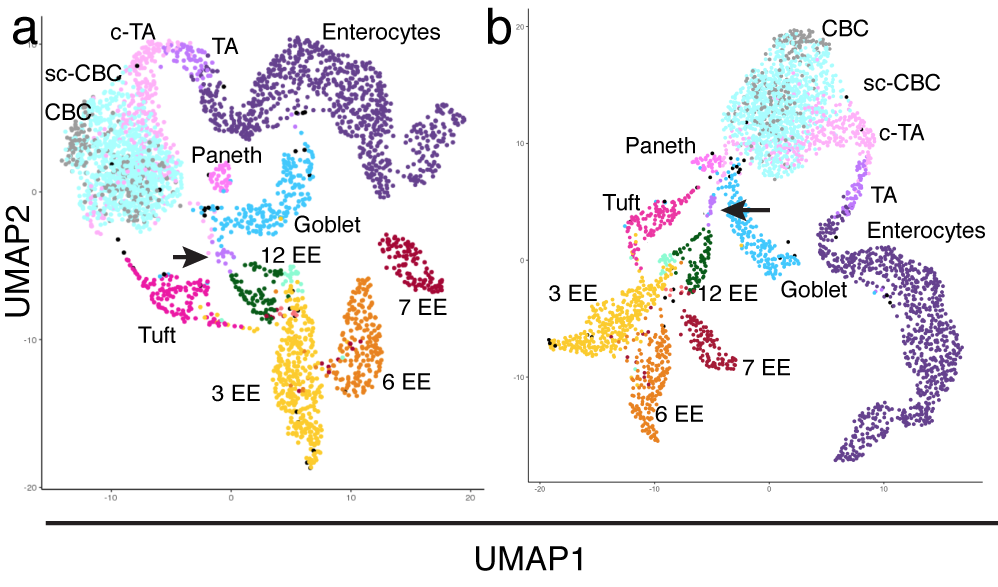

**Fig. S6. UMAP embedding of tSpace distance matrix.** Most developmental branching relationships are retained in 2D UMAP embedding of the dense tSpace distance matrix (**a-b**). UMAP of tSpace matrix with min_dis = 1 **a** Euclidean and **b** Manhattan distance. slEEP cells are marked with an arrow. TA – transit amplifying, CBC – crypt base columnar, sc-CBC slow-cycling CBC, EE – enteroendocrine cells.

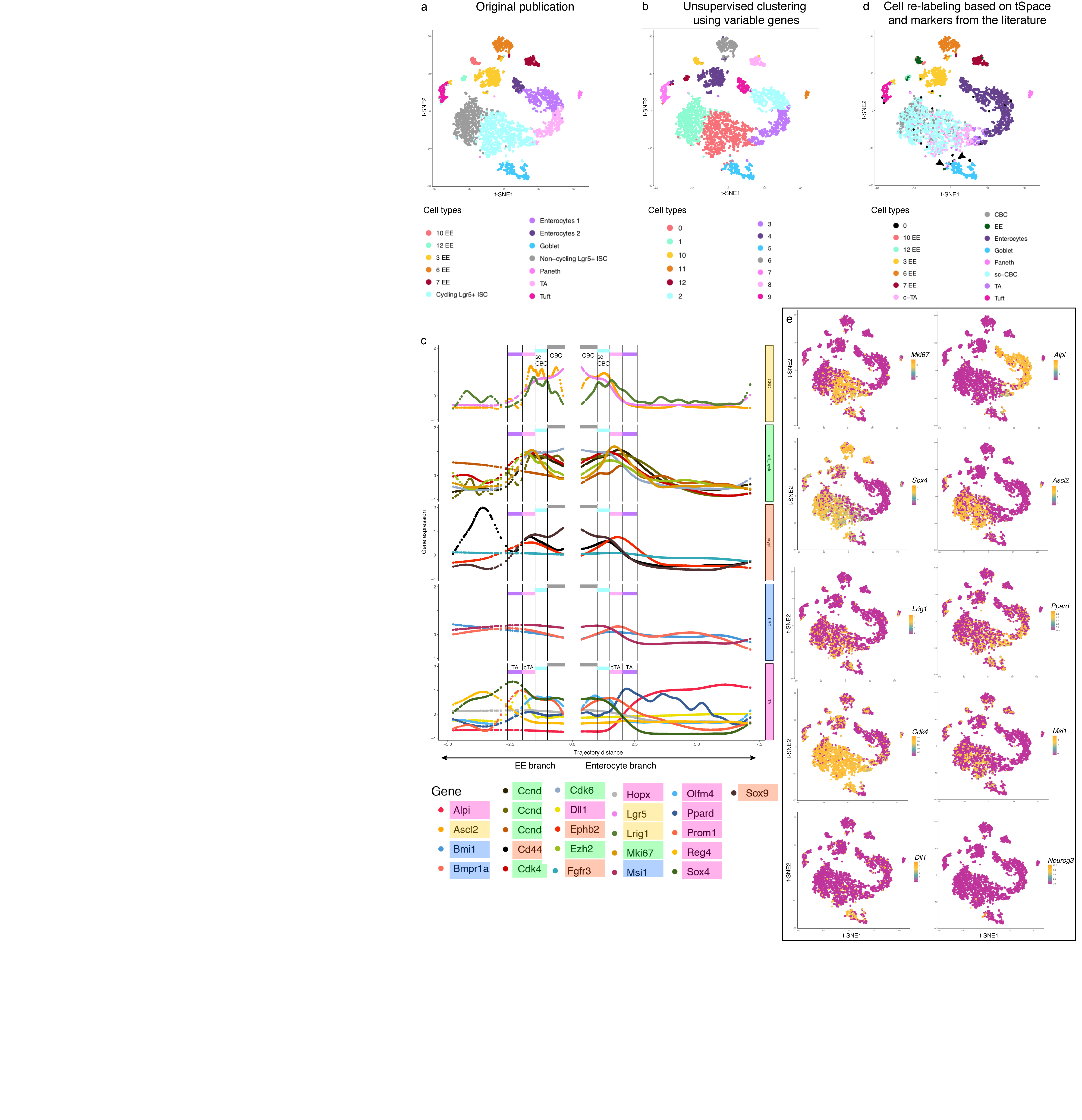
**Fig. S7. Proposed labels for the cell populations within the intestinal crypt based on developmental distance (distances along isolated cell paths from tSpace) and subset markers. a** t-SNE visualization of cell populations, with the original labels^2^. **b** t-SNE visualization of cell populations determined using all variable genes and unsupervised clustering provided in Seurat package. Annotation of subsets is from Yan *et al.* based on unsupervised clustering and association with known intestinal populations. **c** Smoothed expression of markers^5^ that define the crypt zone (orange), cell cycle and proliferation (green), and populations (CBC, TA) associated with the intestinal crypt along isolated trajectories: these markers were used to re-evaluate cell designations, identify putative label retaining cells (LRC, blue), crypt base columnar (CBC, yellow) and transit amplifying (TA, pink) cells. We used peaks of expression of multiple subset-associated genes to define specific cell types. Vertical black lines are suggested boundaries between cell populations. The combination of proliferation markers (green) with CBC and TA cell markers allows separation of slow cycling CBC and cycling TA cells. *Dll1* and *Sox4* are exclusively expressed by TA within the EE branch, while *Alpi* gene and *Ppard* are specific to TA within the enterocyte branch. The cell identities defined provide landmarks for orientation, but tSpace visualization emphasizes that these populations actually exist in the tissue as part of a developmental continuum. **d** t-SNE visualization of the newly tSpace-defined cell populations within the crypt. Arrows point to the slEEP cells, delineated in trajectory space and known to exist in intestinal crypt, but which are not defined by conventional t-SNE, SPADE or unsupervised clustering (Seurat) but rather clustered with goblet cells. Cells labeled with 0 are all cells that did not fall into our vertical gates. **e** Selected markers from **c** panel, shown in a t-SNE map.

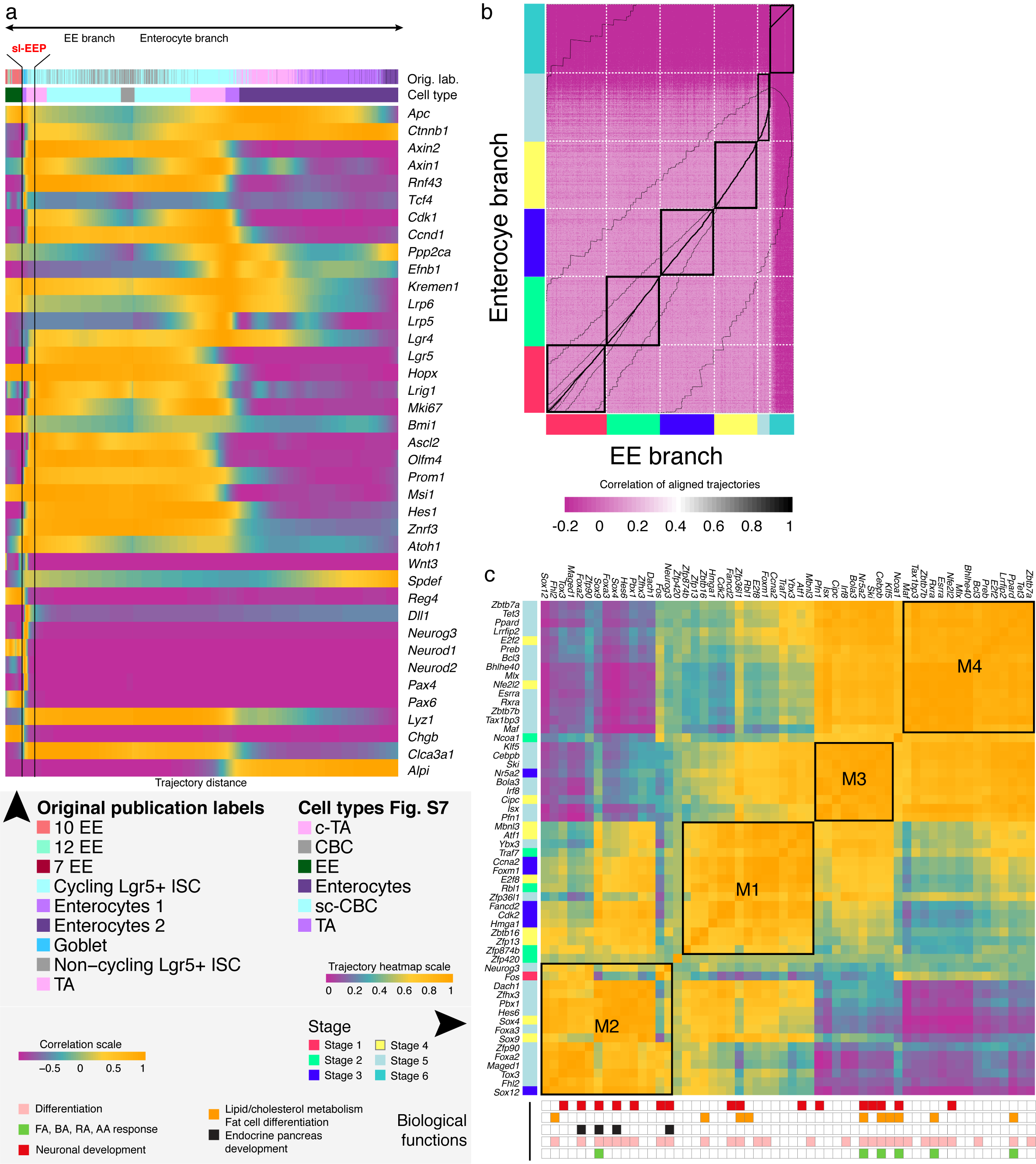

**Fig. S8.** Trajectory analysis allows identification of candidate TF’s and TF modules for enterocyte and enteroendocrine (EE) differentiation. **a** Expression patterns of genes that regulate intestinal crypt development reviewed by Hans Clevers, along the isolated trajectories confirm known biology. Cell labels from original publication, our proposed cell labels and short lived enteroendocrine progenitors (slEEP) are marked along the trajectories. Cell stage (Methods) and cell identities defined here (cell subsets) or in Yan *et al*. (Original labels) are indicated above the heatmap. **b** Heatmap of single cell correlations between two trajectories and alignment cost (thin black contour bars), which is minimal along the thicker black line. Trajectories were aligned with dynamic time warping and subsequently sectioned into 6 segments for further pair-wise comparison (e.g. Stage 1 EE branch vs Stage 1 enterocyte branch). **c** Co-expression analysis of transcription factors identifies 4 modules (M1-M4) specific for intestinal crypt and cell commitment to enterocyte or EE lineages. Stages shown next to the TFs are the earliest stage when TF was significantly changed in one of the compared trajectories. Biological functions of TFs are highlighted in the panel below the correlation heatmap.

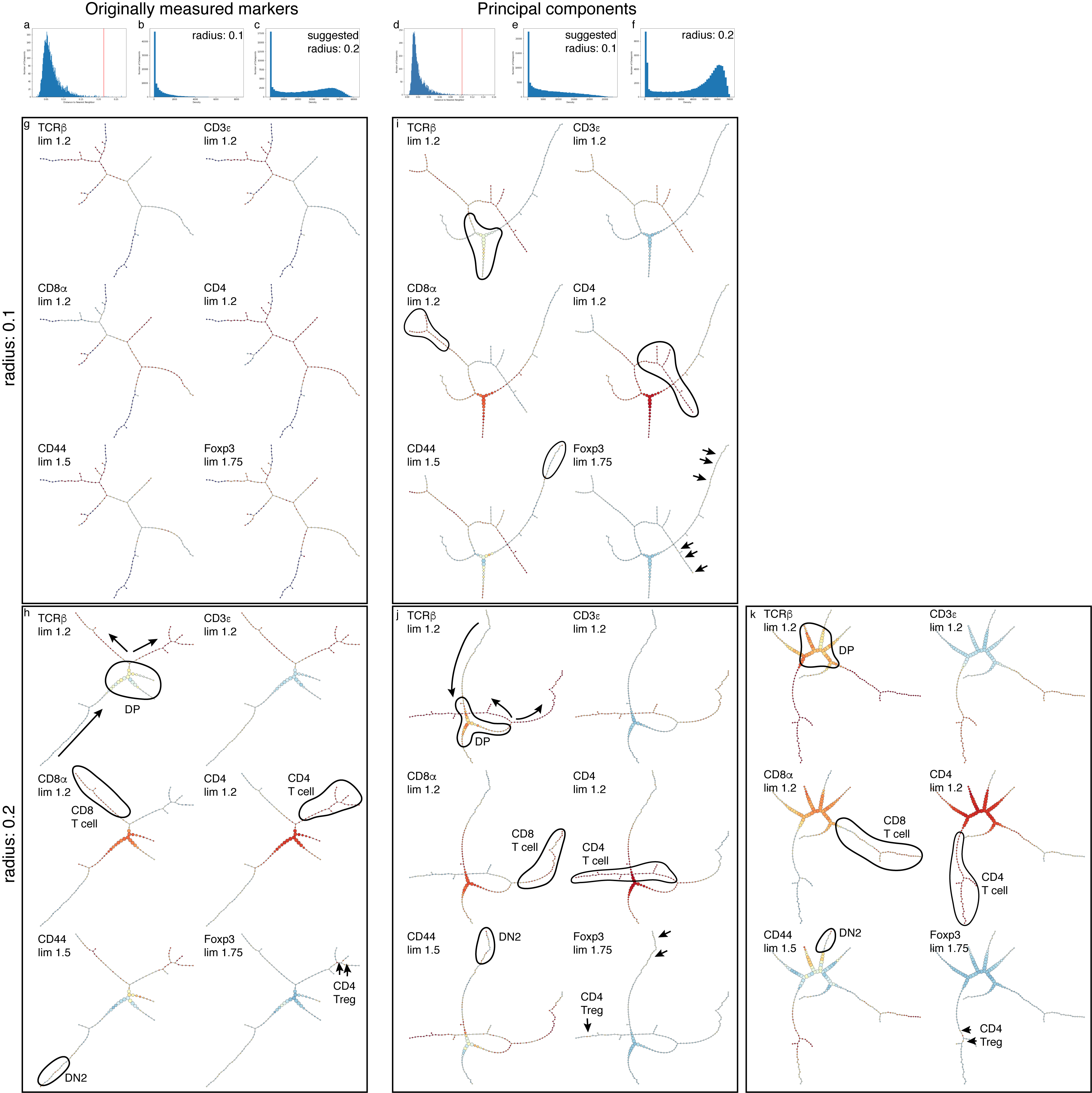
**Fig. S9.** p-Creode analysis of thymocyte development. **p-Creode analysis requires downsampling, is inconsistent and sensitive to choice of parameters.** We ran p-Creode using originally measured markers (g-h) vs. principal components (i-k), which is suggested by p-Creode tutorial. We compared the effect of different values of radius parameter. **a** and **d** suggested radius, **b**, **c,** **e** and **f** cell density graphs for different radius. **g**, **h**, **i**, **j** different p-Creode analysis depended on selection of radius (g & i radius 0.1, h & j/k radius 0.2) and measured markers vs 5 principal components. **k** p-Creode analysis of the same T cell data with the same parameters as in **j**, showing effect of random downsampling on the final output of p-Creode.

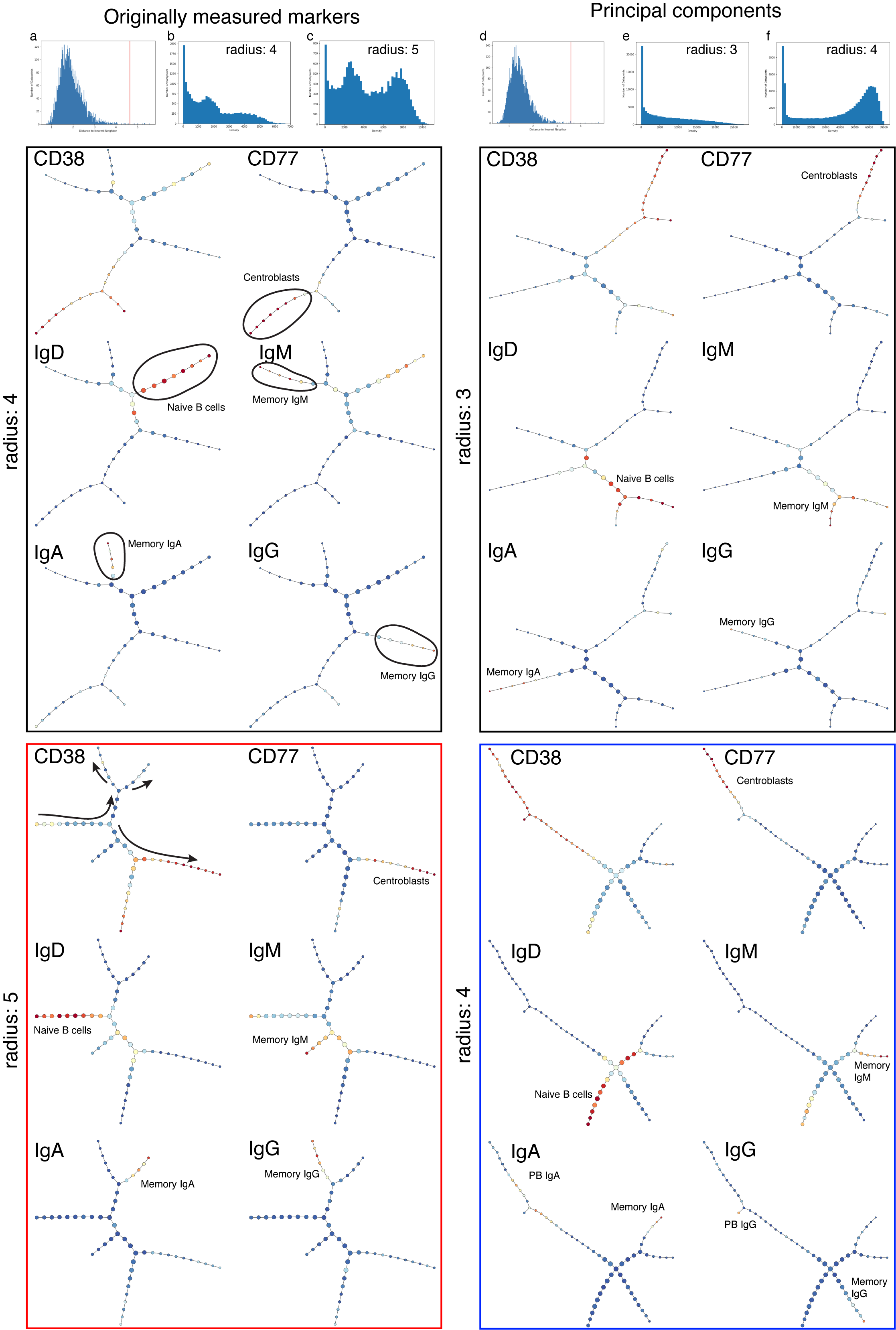
 **Fig. S10.** **p-Creode analysis of tonsil B cells. As for thymocytes, the output is sensitive to choice of parameters. It also fails to differentiate small populations (plasmablasts).** p-Creode of tonsil B cells. We ran p-Creode using originally measured markers (left panels) or principal components (right panels), which is suggested by the p-Creode tutorial; and we compared the effect of different values of the critical radius parameter. Red and blue squares mark analyses that are closest to real biology. **a** and **d** suggested radius, **b**, **c,** **e** and **f** cell density graphs for different radius. Four squares show different p-Creode analysis depended on selection of the radius and on use of measured parameters or 15 principal components. Red and blue squares label closest p-Creode solution to the real biology.

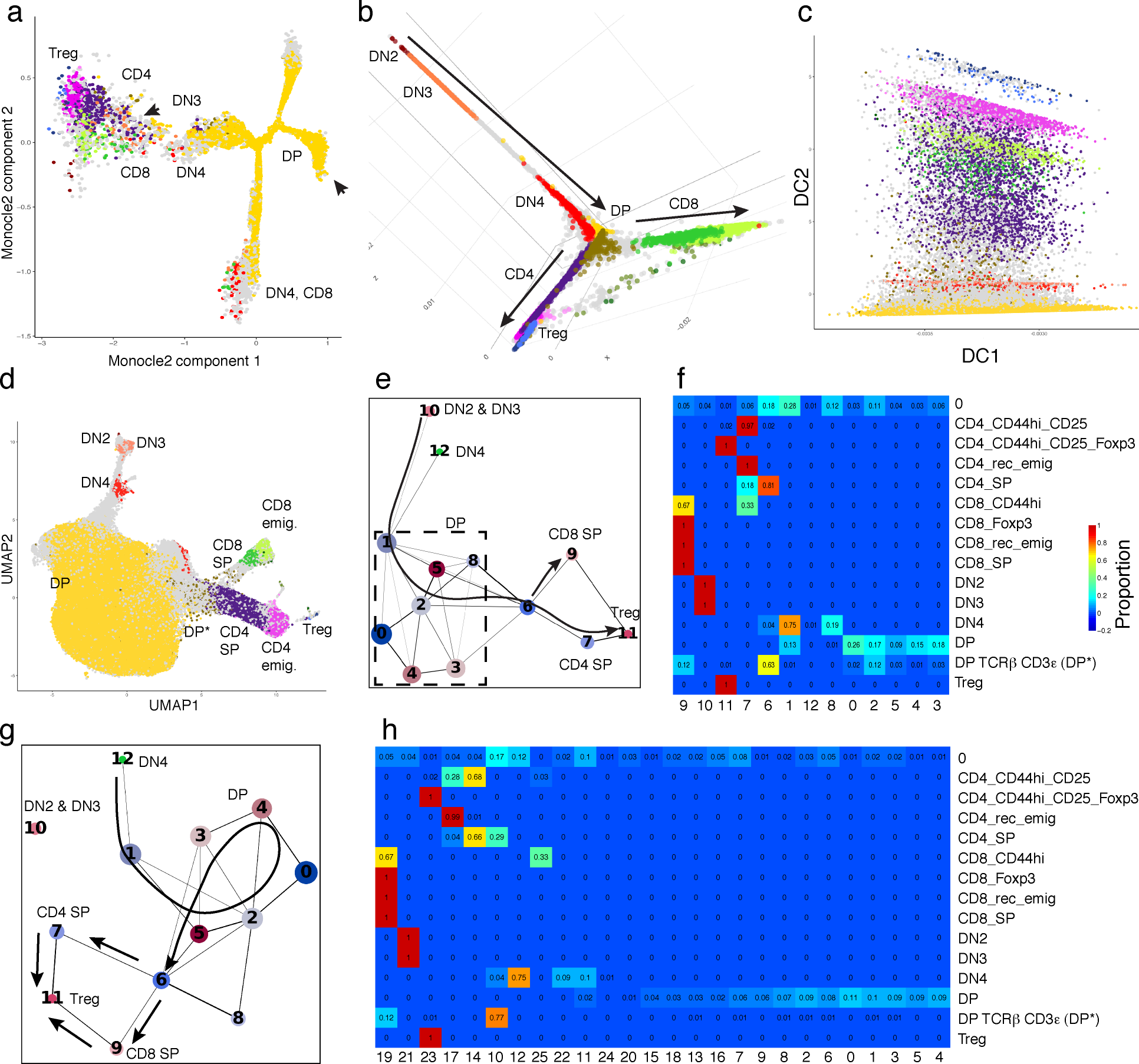

**Fig. S11.** T cell data analyzed using **a** Monocle2, **b-c** diffusion maps, **d** UMAP and **e-h** PAGA. Monocle2 failed to reveal the relatively simple T cell maturation branching. Diffusion maps, if the first diffusion component is ignored (DC1 is shown in panel c) reveals CD4 and CD8 branching as appropriate, however subtle sub-branches (i.e. of Tregs vs conventional CD4 mature T cells) are not prominent. PAGA fails to connect DN2 and DN3 precursors to DN4 and the rest of more mature T cells.

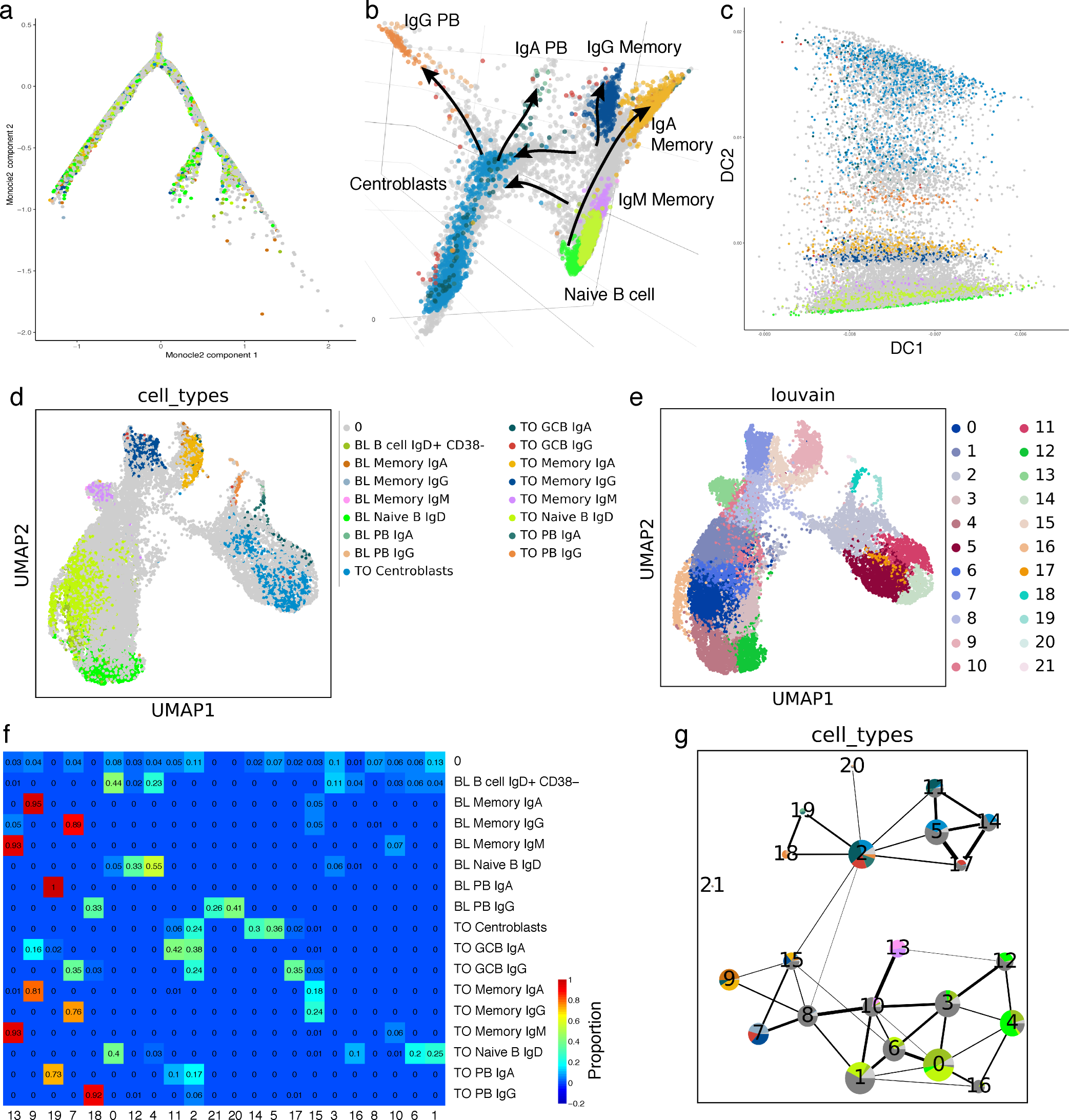

**Fig. S12. B cell data analyzed using a** Monocle2, **b-c** diffusion maps, **d** UMAP, **e-g** PAGA. Monocle2 (a) fails to reveal B cell maturation in tonsil. Diffusion maps (b and c), as in T cell data if the first diffusion component is ignored, reveals central developmental paths in B cell development but positions IgM memory cells along the core path from naïve cells to germinal center cells (GCC); but obscures links from GC to memory cells that are revealed by intermediates in tSpace. UMAP (d) clusters cells obscuring intermediate populations and potential alternative differentiation sequences. PAGA (e-g) connects naïve cells with germinal center (GC) B cells through IgG and IgA memory B cells, without direct connection between naïve and GC.

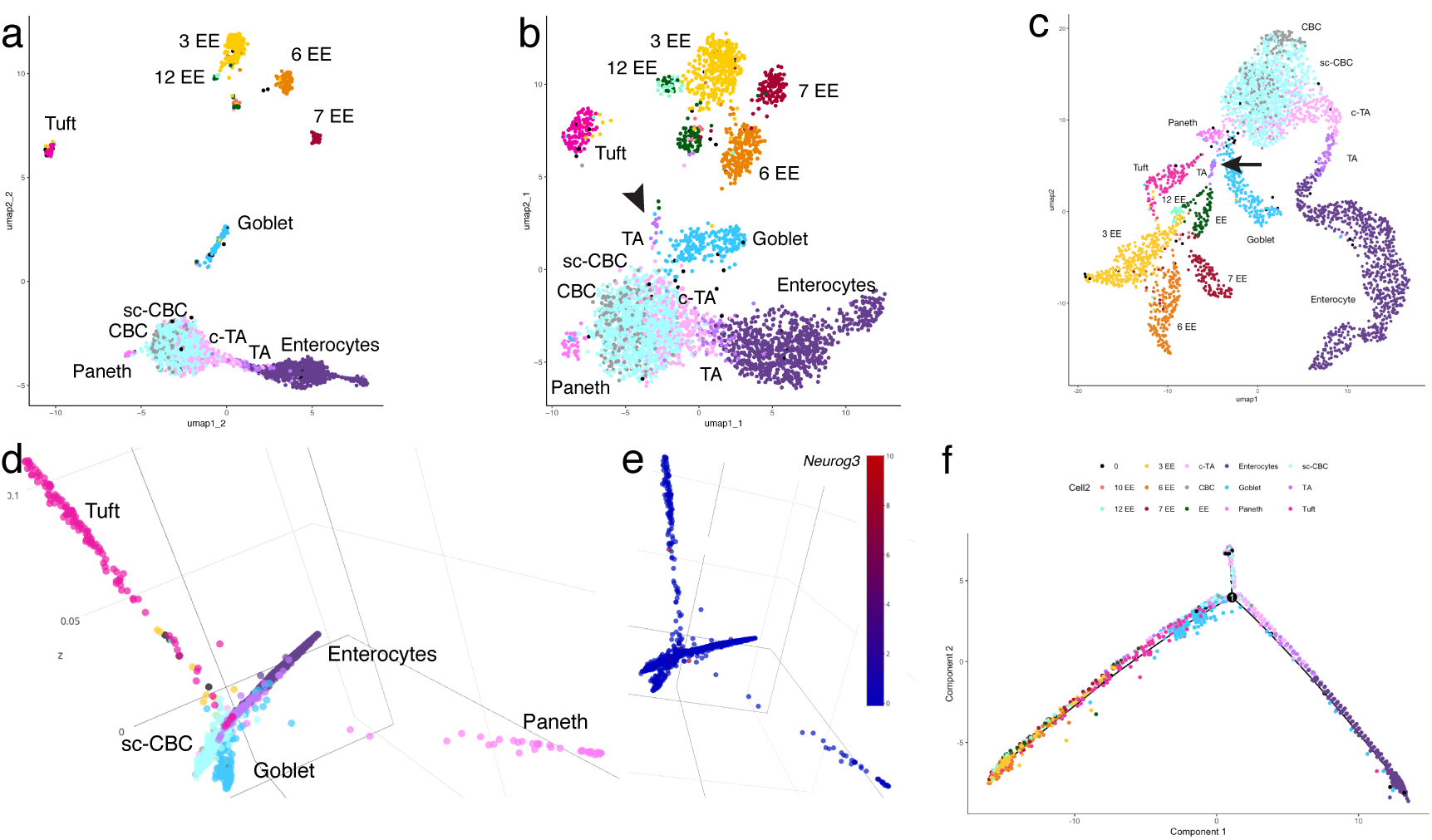

**Fig. S13. Diffusion map and UMAP analysis of mouse small intestine.** UMAP of gene expression with Euclidean distance and different min_dis parameters **a** 0.1, **b** 1. UMAP on variable genes, even with the “loosest” setting for min_dis (b) fails to connect slEEP cells to the rest of secretory subsets. **c** UMAP embedding of cell trajectory profiles (tSpace matrix), in contrast to UMAP of gene expression profiles (b), distinguishes sIEEP cells and correctly positions them on a path from stem cells to EE populations, which are well separated in the trajectory space UMAP projection. UMAP embedding, for visualization, of the trajectory space matrix (here and in Fig. S6) and PCA embedding of tSpace (as shown in text Fig. 3a) outperform all of the methods we evaluated. Arrow marks position of slEEP cells. **d** Diffusion maps with all cell types. Diffusion map branches Paneth, goblet, tuft and enterocytes, while the rest of secretory subsets are not represented, and it fails to show connection between crypt base columnar cells and the rest of differentiated cells, suggesting that diffusion maps perform well in determination of differentiated cells, however **e** *Neurog3* gene expressing cells, described in the manuscript as slEEP cells are lost in the cloud of early stem cells. **f** Monocle2 reveals only two major branches absorptive and secretory, without any subtleties, and sub-branches of the secretory branch are missing.

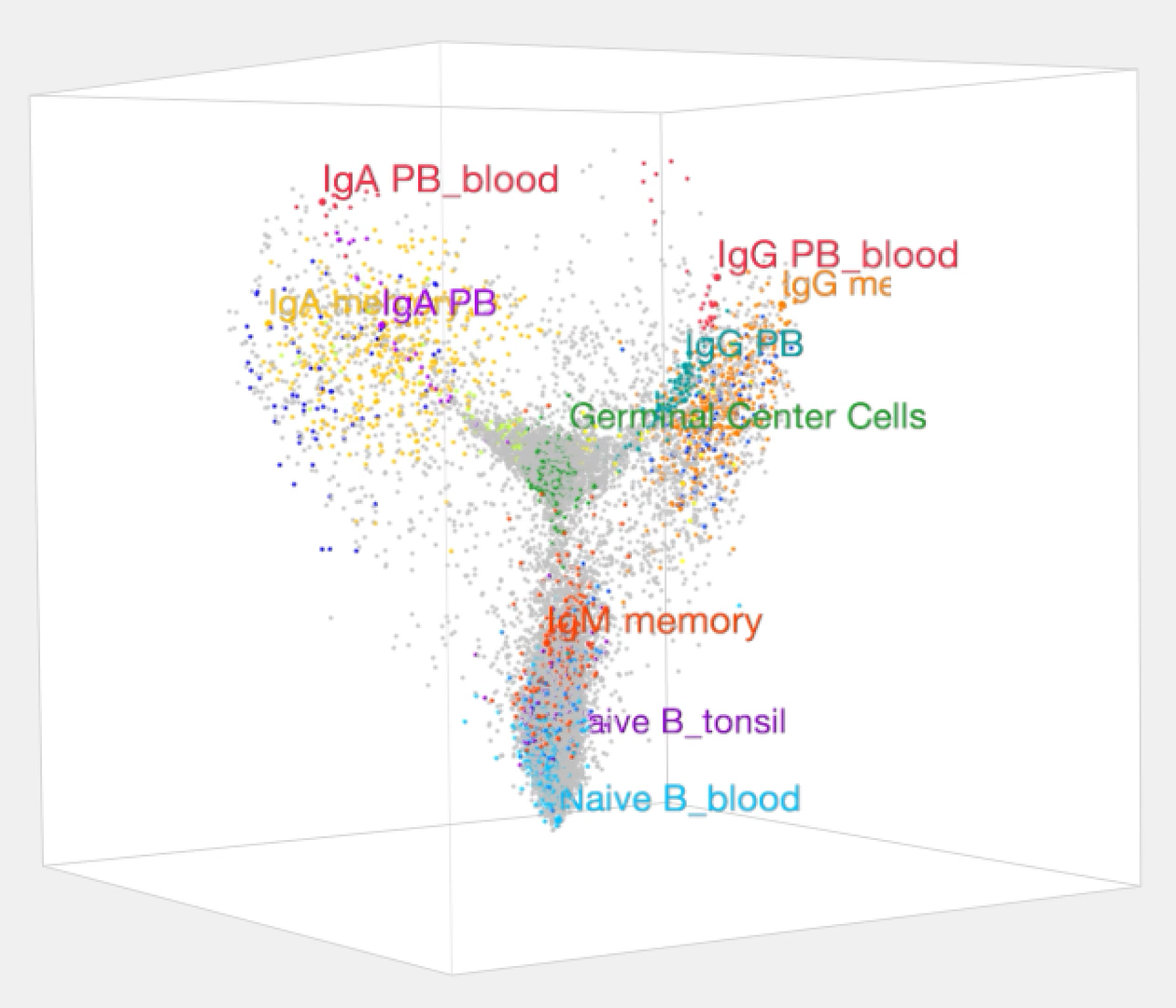

**Supplementary Movie 1. 3D embedding of tSpace principal components reveals B cell differentiation in tonsils and alignment with B cells in blood.** Rotation of the 3D imbedding facilitates visualization of naïve B cell trunk, germinal center, memory B cell and plasmablast populations and their relationships, and the position of intermediate populations. Naïve blood and tonsil B cells, and memory IgA and IgG B cell pools exchange between blood and tonsil, and are intermixed in the projection. In contrast developing plasmablasts mature in the tonsil and migrate into the blood: alignment in trajectory space links terminal tonsil plasmablast development directly to their blood counterparts.

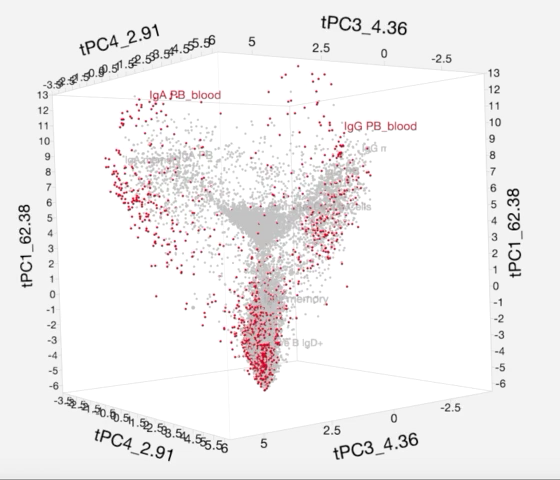

**Supplementary Movie 2. Exchange of blood naive and memory but not GC B cells seen in trajectory space.**
